# Cross-species examination of X-chromosome inactivation highlights domains of escape from silencing

**DOI:** 10.1101/2020.12.04.412197

**Authors:** Bradley P Balaton, Oriol Fornes, Wyeth W Wasserman, Carolyn J Brown

## Abstract

**Background:** X-chromosome inactivation (XCI) in eutherian mammals is the epigenetic inactivation of one of the two X chromosomes in XX females in order to compensate for dosage differences with XY males. Not all genes are inactivated, and the proportion escaping from inactivation varies between human and mouse (the two species that have been extensively studied).

**Results:** We used DNA methylation to predict the XCI status of X-linked genes with CpG islands across 12 different species: human, chimp, bonobo, gorilla, orangutan, mouse, cow, sheep, goat, pig, horse and dog. We determined the XCI status of 342 CpG islands on average per species, with most species having 80-90% of genes subject to XCI. Mouse was an outlier, with a higher proportion of genes subject to XCI than found in other species. Sixteen genes were found to have discordant X-chromosome inactivation statuses across multiple species, with five of these showing primate-specific escape from XCI. These discordant genes tended to cluster together within the X chromosome, along with genes with similar patterns of escape from XCI. CTCF-binding, ATAC-seq signal and LTR repeats were enriched at genes escaping XCI when compared to genes subject to XCI; however, enrichment was only observed in three or four of the species tested. LINE and DNA repeats showed enrichment around subject genes, but again not in a consistent subset of species.

**Conclusions:** In this study we determined XCI status across 12 species, showing mouse to be an outlier with few genes that escape inactivation. Inactivation status is largely conserved across species. The clustering of genes that change XCI status across species implicates a domain-level control. In contrast, the relatively consistent, but not universal correlation of inactivation status with enrichment of repetitive elements or CTCF binding at promoters demonstrates gene-based influences on inactivation state. This study broadens enrichment analysis of regulatory elements to species beyond human and mouse.

## Background

Human and mouse differ in both the initiation and completeness of X-chromosome inactivation (XCI) [1,2]. In contrast to human, mouse has imprinted XCI early in development, which is maintained in extraembryonic (placental) tissues [3–5]. In placenta, rat [6] and vole [7] also have imprinted XCI while horse/donkey hybrids [8] and pig [9] have random XCI. The story is unclear in cow, where both random [10] and imprinted [11] XCI have been reported. At the blastocyst stage, human as well as rabbit express XIST (the RNA that initiates the silencing cascade) from both alleles, while mouse has exclusively paternal Xist expression [1]. Cow has been observed to upregulate XIST at a similar stage to human and rabbit [12]. Human and rabbit also showed later inactivation timing than mouse [1]. See [13] for a review of XCI across species.

Not all genes are subject to XCI, and here again, there is a substantial difference between human and mouse. Escape from XCI is generally defined as having an inactive X (Xi) expression of at least 10% of active X (Xa) expression [14]. Around 12% of X chromosome genes are escaping XCI in human [15], while in mouse the proportion of genes escaping from XCI is only 3-7% [16]. In human, an additional 15% of genes variably escape from XCI, differing in their XCI status between different tissues, populations, individuals or studies [15,17]. Large-scale studies have not been reported in species outside of human and mouse, and the studies in mouse generally report only on the genes escaping from XCI. The variation between species highlights the importance of studying XCI across a range of species; particularly as the most common model organism, mouse, appears quite different from human.

There are various methods to examine the XCI status of genes, with the above numbers being determined using a combination of allelic expression and DNA methylation (DNAme). Additional methods to assess XCI status are reviewed in [18]. For allelic expression to be used to examine XCI escape status, the samples analyzed must be skewed so that the majority of cells in the sample have the same Xi. Skewing of XCI >90% occurs infrequently in human, but at elevated incidence in blood [19] and cancer due to its monoclonal origin [20]. Cell lines that have undergone clonal selection or which are skewed due to X-linked diseases have also been used [14]. Mouse lines with the gene that controls initiation of XCI, *Xist*, knocked out on one allele exclusively inactivate the X chromosome with functional *Xist* [16]; and selectable markers such as fluorescent proteins can also be inserted on one of the X chromosomes in order to select for cell populations with a consistent Xa [21]. Trophoblast cells in mouse have imprinted XCI, and have also been used to determine XCI status [22]. Overall, the requirement for skewing of XCI dramatically limits the datasets that can be used to analyze escape from XCI using allelic expression.

DNAme-based analyses circumvent this challenge. DNAme of CpG islands at promoters is strongly predictive of XCI escape status [23]. CpG islands are regions of at least 200 bp with high GC content and limited depletion of CG dinucleotides, and are often associated with the promoters of genes, particularly house-keeping genes [24]. Males have low DNAme of promoter CpG islands on the X chromosome, while females, with one Xa and one Xi, will have one relatively unmethylated chromosome and one methylated chromosome, for an average methylation level around 50%. DNAme in gene bodies differs between genes escaping from and subject to XCI, but these differences are subtler and may be tissue-specific [23,25–27]).

Knowing the XCI status of genes is important, as genes that escape from XCI often have sex-biased expression, being higher in males if a gametolog is also present on the Y, and higher in females if not [17]. Furthermore, having two active copies of a gene has been argued to protect females from cancers as both copies will need to be mutated in order to have loss of function [28]. In individual species, knowing which genes escape from XCI will be useful for mapping the effect of X-linked genes to various traits, and understanding XCI within a species is important for genomic selection strategies in breeding for agriculture [29]. Additionally, the knowledge of which genes escape from XCI across species can further our understanding of the underlying mechanism allowing some genes to escape XCI and give insight into the evolutionary development of XCI.

Here, we compared the XCI status of human and mouse, first examining allelic expression and DNAme in human and mouse to establish robust thresholds of DNAme as an indicator of XCI. We then used DNAme data across two separate groups, one of nine different mammalian species, and one of five different primate species, to examine conservation of XCI escape status across species. Finally, we performed an analysis testing elements previously seen enriched at genes with various XCI statuses (repetitive elements, CTCF and ATAC-seq) for enrichment with our XCI status calls across species.

## Results

### XCI status calls from allelic expression

To obtain DNAme thresholds separating genes escaping XCI from genes subject to XCI, we first needed to establish which genes were escaping versus subject to XCI using allelic expression data. Allelic expression data requires skewed Xi choice and thus was only available for two species: human and mouse (**Figure 1, S1, S2)**. Expression-based XCI status calls were determined using a binomial model as previously described [16], with genes having an Xi/Xa expression ratio significantly over 0.1 being called as escaping XCI and those with Xi/Xa significantly under 0.1 being called as subject to XCI. For human, our skewed samples were from cancer-related samples and we identified 44 genes escaping XCI, 262 genes subject to XCI and 21 genes variably escaping from XCI (**Table S1)**. The majority of these XCI status calls agreed with previous studies, with discordance for only 53 genes, (17% of genes with an XCI status call in both), 39 of which were reported to variably escape from XCI here or previously [15]. We attribute the low number of genes variably escaping in our current study to the limited number of samples available and the frequency of informative, heterozygous SNPs per sample, resulting in a mean of 3.5 informative samples per gene. With more samples, we would expect to observe more variably escaping genes.

**Figure 1:**
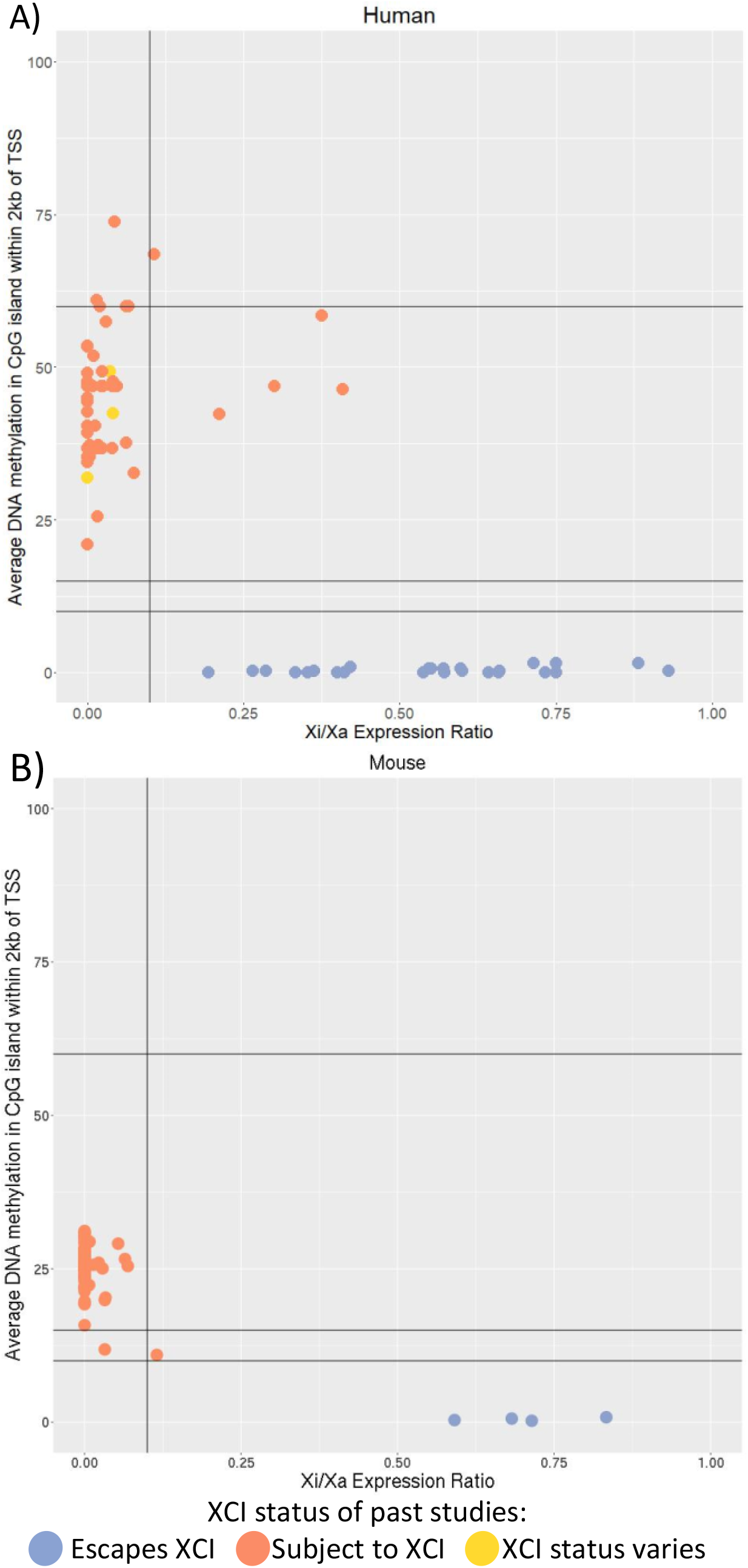
Using Xi/Xa expression ratio to establish thresholds of DNAme for XCI status calls. Two species are featured: human (A) and mouse (B). Each point is a SNP with Xi/Xa expression data, matched to the closest CpG island within 2kb of the closest TSS (accounting for which splice variants would include the SNP) in order to have matched DNAme values. Lines are drawn at 0.1 Xi/Xa expression and at 10, 15 and 60% DNAme as they were used as thresholds to call XCI escape status subsequently. Points are colored based on their XCI status calls. For human, previously published XCI status calls were used [15], while in mouse, which did not have studies calling genes as subject to XCI, they were colored based on their Xi/Xa expression-based XCI status calls featured here. Genes in the pseudoautosomal region, which matches to the Y chromosome, were filtered out.

In mouse we classified 16 genes as escaping XCI, 662 genes subject to XCI and 10 genes variably escaping from XCI (**Table S2**). We used three different mouse expression datasets (Keown *et al*., Berletch *et al*. and Wu *et al*.) and results were 97%, 90% and 87% concordant when datasets were compared with each other [16,21,26]. Most of the discordance in our results arise from identifying more genes variably escaping in the Wu dataset than the other two datasets. Additionally, our use of a threshold of 0.1 rather than 0 to call escape from XCI and the inclusion of a variable escape category resulted in more discordant calls relative to those assigned by Berletch [16]. Figure 1 shows a clear DNAme difference between genes with an Xi/Xa expression ratio under this 0.1 threshold and genes with an Xi/Xa expression ratio over the threshold.

### XCI status calls from DNAme

DNAme data has also been used to call XCI status [23], and is now available from a number of species where expression in individuals with skewed Xi choice is not available. Our search of GEO [30] for DNAme data across eutherian species found datasets with females for 12 different species: human, chimp, bonobo, gorilla, orangutan, mouse, cow, sheep, pig, horse, goat and dog (**Table S3**). Most of the datasets used whole genome bisulfite sequencing (WGBS), while horse was limited to a reduced representation bisulfite sequencing (RRBS) dataset and many of the primates and dog were processed on the Illumina Infinium Human Methylation450 BeadChip array (450k array), with probes that didn’t map well to the species in question being filtered out by the source publications. Plotting male versus female DNAme at promoter CpG islands on the X chromosome showed similar trends across species (**Figure S3**) with a cluster of sites with less than 10% methylation in both, the bulk of sites showing higher female and low male methylation, and the cluster that is over 70% methylated in both sexes being under-represented on the array data. There are some differences in the amount of male hemi-methylated islands and the female DNAme average across species, which could be due to differences across species or due to the different tissues and methods of assessing DNAme used.

DNAme levels for human and mouse were compared to Xi/Xa expression in order to establish thresholds of DNAme for calling escape from XCI (**Figure 1, S1, S2**). There was good correlation between XCI status calls made using Xi/Xa expression and DNAme with a 10% DNAme threshold. An uncallable zone between 10-15% DNAme was added to lower the chance of miscalling genes, as most discordancies between Xi/Xa expression-based calls and DNAme- based calls had DNAme levels in this range. DNAme at genes subject to XCI was lower than expected if the Xi was completely hypermethylated, with an average DNAme of 38% and 27% in human and mouse, respectively (**Table 1**). This shows that the DNAme on the Xi is not complete at these CpG islands. Looking at autosomal imprinted genes, the expected 50% DNAme ratio was found, demonstrating that lower methylation is not a problem inherent with this analysis or datasets, rather it reflects the DNAme levels of the Xi (**Figure S4**).

**Table 1:** Mean DNAme for genes subject to XCI per dataset.

| Species | Data Type | Average DNAm |
| --- | --- | --- |
| Human | WGBS | 38% |
|  | 450k array | 41% |
| Chimp | WGBS | 35% |
|  | 450k array | 41% |
| Bonobo | 450k array | 38% |
| Gorilla | 450k array | 39% |
| Orangutan | 450k array | 39% |
| Mouse | WGBS | 27% |
| Cow | WGBS | 37% |
| Sheep | WGBS | 31% |
| Goat | WGBS | 33% |
| Pig | WGBS | 38% |
| Horse | RRBS | 37% |
| Dog | 450k array | 39% |

*The mean DNAm of CpG islands at genes found subject to XCI was calculated per dataset.*

Applying our DNAme thresholds across species to make XCI status calls generated between 26 and 567 XCI status calls per species, with a median of 342 calls per species **(Table S1, S2)**. Most species had 80-90% of genes identified as subject to XCI by DNAme (**Figure 2**), while mouse had 95% of genes subject to XCI and horse only had 76% of genes subject to XCI. The decreased number of genes subject to XCI in horse may be due to the data being generated using RRBS, which provides sparser data and, unlike 450k data, the sparse CpGs assessed are not the same across samples. In other species the average DNAme at genes subject to XCI ranged from 31% in sheep to 41% in the chimp 450k array data. The 450k array data tended to have higher DNAme than WGBS data, with values between 38 and 41%. Comparison between human and chimp WGBS and 450k array data at the same genes showed that the WGBS and 450k data differ in DNAme levels, with R^2^ values of 0.04 in chimp and 0.59 in human (**Figure S5**). Differences may be due to having more CpG sites averaged in the WGBS data. XCI status calls made using our DNAme thresholds were consistent however, so we did not discard the 450k array datasets. Of the genes that had XCI status calls from both methods, 98% of human genes had the same calls as did 92% of chimp genes. Most of the genes discordant between methods were hypermethylated in one method, and therefore were not given an XCI status call in that analysis.

**Figure 2:**
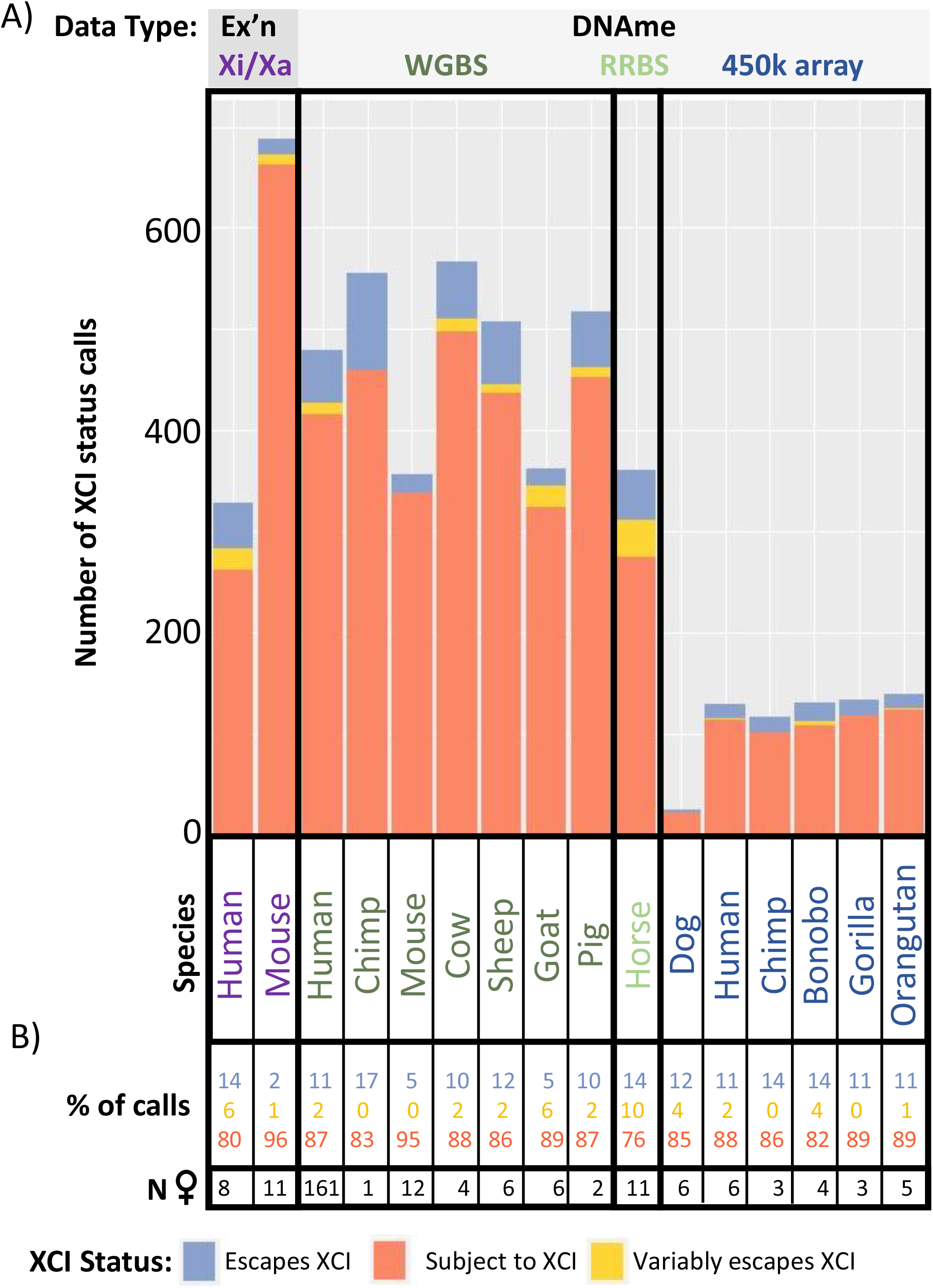
The number and type of XCI status calls per species. The number of XCI status calls per dataset (A) and the percentage of calls with each XCI status per dataset (B) are shown. Datasets (columns) were sorted by technique used to generate the data. Species names are colored by the type of data used to generate XCI status calls.

Horse had elevated numbers of variably escaping genes (10%), which was close to that seen previously in human, while other species (including human) only had 0-5% of genes found variably escaping from XCI. The variation in proportion of variable escape genes seen here could be due to low sample size (in everything except human WGBS), or from our methods of calling variable escape genes being more stringent than previous studies. We required at least 33% of informative samples to have each XCI status before calling a gene as variably escaping from XCI, similar to the initial survey of human XCI status by Carrel and Willard [14]. Reducing this requirement to only 10% of samples increased the number of variably escaping genes found in human to 63 - almost a quarter of informative genes. These include 37 new genes called which did not have enough informative samples to be called as escaping or subject to XCI with our initial thresholds, as well as 15 genes which changed from an initial call of escaping XCI (12 genes) or subject to XCI (three genes). Although this lower threshold called more genes, we used our 33% threshold of variable escape calls for subsequent studies as we wished to focus on genes that we were confident changed their XCI status between species, rather than differing levels of variable escape from XCI.

Overall, we saw that calls of XCI status using DNAme agreed well with those made using allelic expression, and provided an opportunity to examine XCI across multiple species. While WGBS resulted in the most XCI status calls, 450k DNAme-based calls were generally concordant. These studies showed an average of 11% of genes escaping from XCI across 12 different species, with mouse being an outlier with only 5% of genes escaping from XCI.

### Conservation of XCI status calls across species

XCI status calls per gene were compared across species, focusing on genes that were informative in 4+ species. We observed 267 genes being completely conserved across all informative species, with only eight of these genes escaping from XCI and the rest being subject to XCI. Of the eight conserved XCI escapees, two (*DDX3X* and *KDM6A*) have Y homologues across eutherian mammals [31], five have Y pseudogenes in human (*ARSD, STS, PNPLA4, EIF2S3* and *MED14*) [32], and one has no known Y homology (*CTPS2*) (**Figure 3A**). To avoid biasing the analysis with the more conserved primates, the species were grouped into two groups: primates with 450k array data, and other datasets (including the human and chimp WGBS data). A clear difference in conservation of status was seen between these two groups, with 97% of genes having completely conserved XCI status across primates, while only 75% of genes had conserved XCI status across all mammals **(Table S1)**. Of the genes which were usually subject to XCI (>75% of informative species subject to XCI), 79% of these had all informative species subject to XCI. Genes that usually escaped from XCI were less concordant, with only 61% of these genes having entirely conserved XCI status across all informative species. A similar trend was seen in the all primates group.

**Figure 3:**
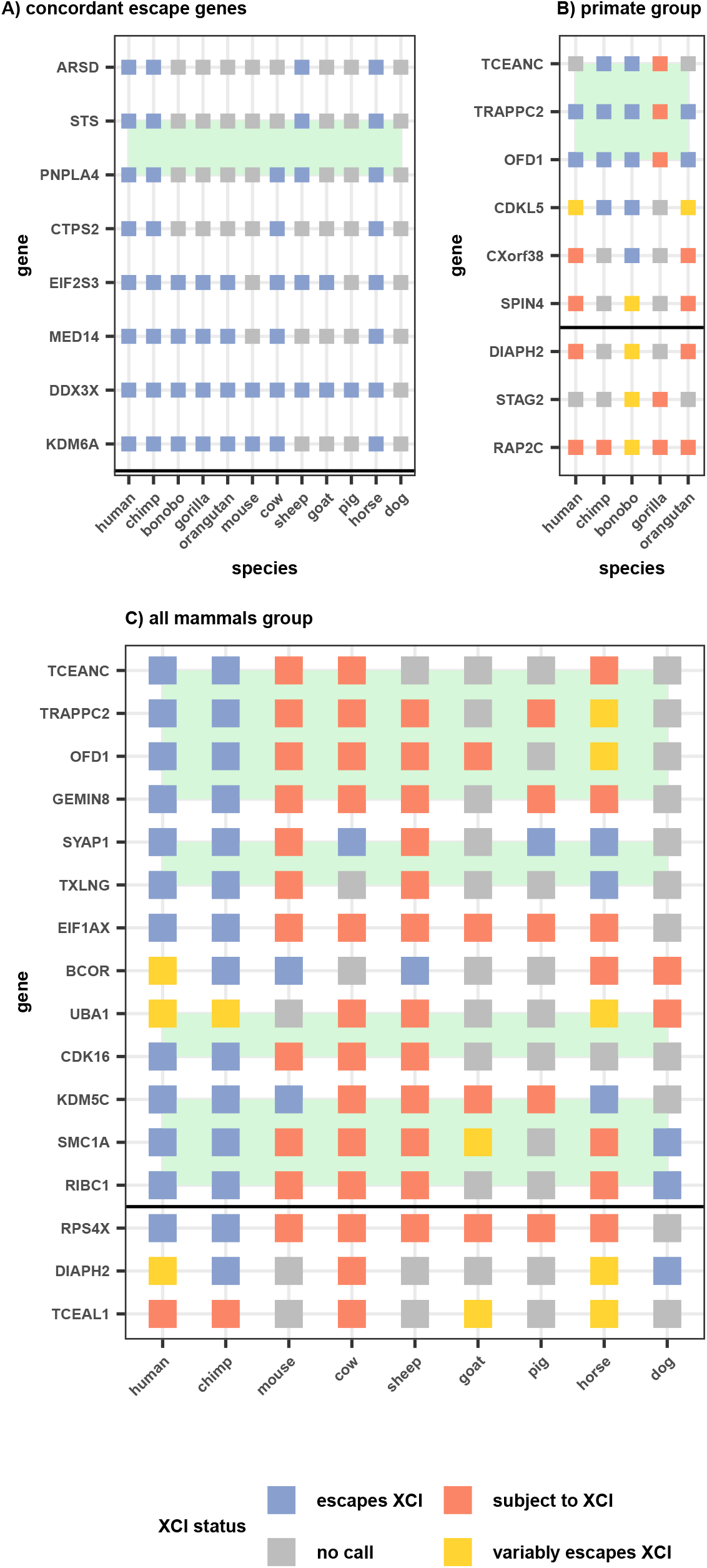
Concordant and discordant escape genes across species. Eight genes escape XCI in all informative species (A), while 259 genes were subject to XCI in all informative species (not shown). Discordant genes in two different groups of species were examined, only primates (B) and all mammals (C, limited to only 2 primate species). The intersection of a gene and species is colored based on that gene’s XCI status call in that species. Genes that did not have an XCI status call in a species are colored grey. Only escape genes informative in at least 4+ species were selected for A. Genes were selected for B if they had at least one discordant primate species while genes in C required two XCI statuses with two or more species. To match best across species within groups, 450k array data was prioritized in B and WGBS data was prioritized in C. Genes are organized based on their position on the human X chromosome with a horizontal black line denoting the centromere. Green boxes highlight domains of adjacent genes with similar changes to XCI statuses across species.

There were 16 genes that varied frequently (2+ species escaping XCI and 2+ species subject to XCI) in the all mammals group and none that varied greatly across primates, again showing the higher similarity in XCI status across closely related species **(Figure 3)**. Of these 16 genes, four showed primate-specific escape from XCI (*RPS4X, CDK16, EIF1AX* and *GEMIN8*) and one showed artiodactyla-specific (cow, sheep, goat, pig) XCI (*KDM5C*). The pattern of conservation of the other genes variably escaping across species did not match any phylogenetic patterns. The primate-specific escape genes *RPS4X* and *EIF1AX* have been shown to have primate-specific retention of their Y homolog while *KDM5C*, the gene that is subject to XCI only in artiodactyla has lost its Y homolog in bulls, while retaining it in mouse and primates [31]. We show the WGBS data surrounding the CpG island at the transcription start site (TSS) of the ubiquitous escape gene *KDM6A*, the artiodactyla-specific subject gene *KDM5C* and the primate-specific escape gene *RPS4X* (**Figure 4**).

**Figure 4:**
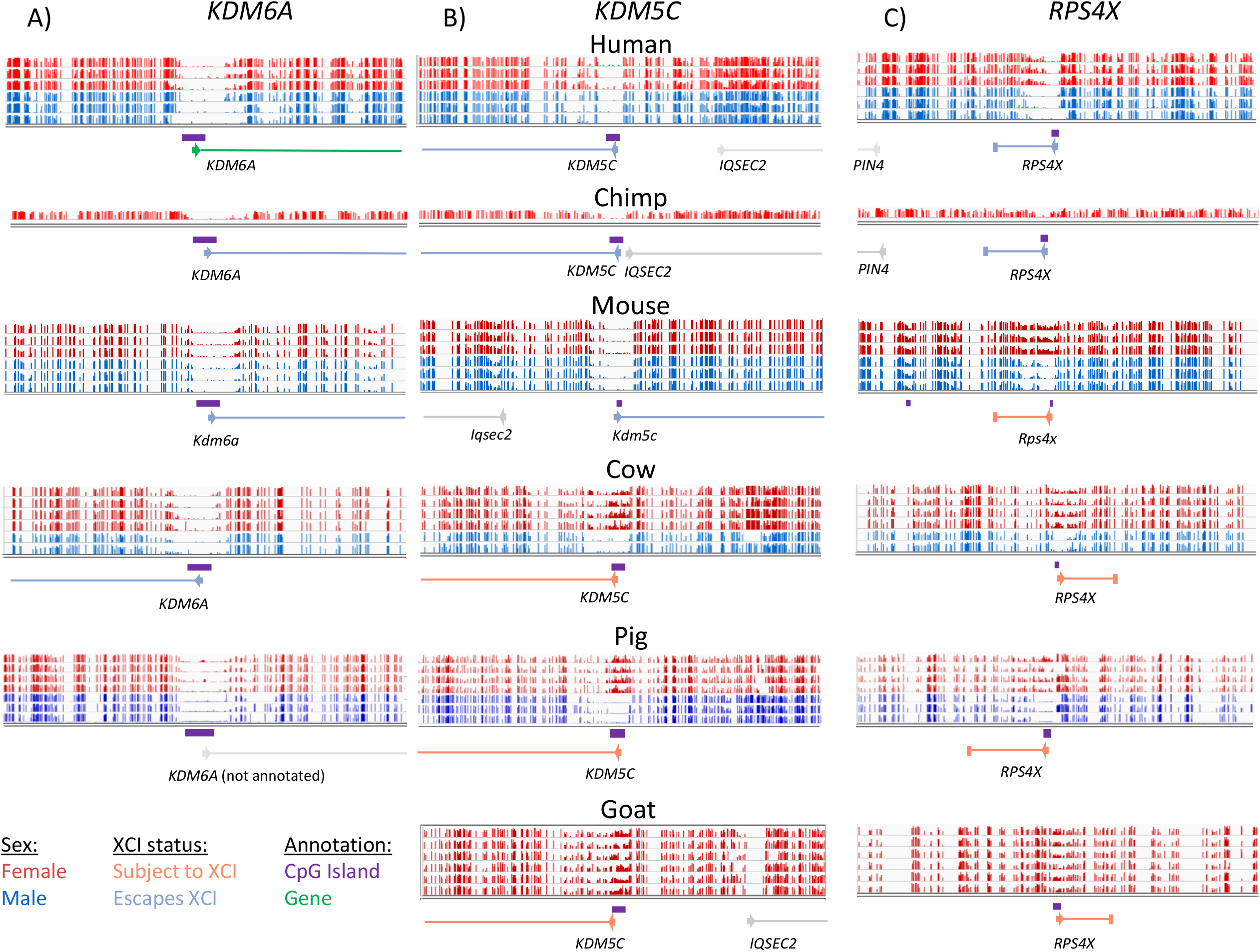
Featured genes compared across species. Male and female DNAme values are graphed by gene and dataset. *KDM6A* is featured as it is concordantly escaping across species (A). *KDM5C* is featured because it is known to escape XCI across species but is here shown to be subject to XCI in artiodactyla (cow, sheep, pig and goat) (B). *RPS4X* is featured because it is a well known primate-specific escape gene (C). Male methylation is shown in blue and female in red. Annotated CpG islands are shown under the methylation data in orange. Genes are shown in green with arrows at the TSS pointing in the direction of transcription. All of the methylation data shown is from WGBS. Pig did not have *KDM6A* annotated, but predictions from other species show it located at this CpG island. Goat did not have a CpG island or hypomethylated region at the annotated *KDM6A*.

*CDKL5* was the only gene seen to have more than one discordant species in primates (**Figure 3B**), being subject to XCI in the human WGBS data, variable in orangutan and the human 450k array data and escaping in chimp and bonobo. In gorilla, *CDKL5* appeared subject to XCI, but half of the data was in the uncallable region between 10 and 15% DNAme so it was not called as subject to XCI. Other genes had only one species of primates discordant from the rest, usually gorilla or bonobo.

*UBA1* was particularly interesting as it has been shown previously in human to have two different TSSs with differing XCI statuses [33]. This pattern of multiple TSSs with differing XCI status was seen also in chimp and horse (although data is sparse in horse) **(Figure 5)**. In cow, the upstream TSS and CpG island are not annotated, but the region homologous to the human upstream TSS showed a DNAme pattern consistent with a promoter subject to XCI, and in pig the CpG islands are annotated but the gene is not. Similarly, in mouse both TSSs (which are annotated but lack CpG island definition) had female-specific DNAme. Mouse has been shown to have fewer CpG islands than human, with CpG island loss from the ancestral genome being four times as high in mouse as human [34]. The island is still large enough to see hypomethylation on the Xa so the cutoff for minimum island size may be too high in some species. Overall, the alternative TSSs are conserved across species; however, the XCI status of the downstream TSS changes from escaping from XCI in human, chimp and horse to being subject to XCI in mouse and cow. Examining TSS usage in the other genes featured in Figure 3C, we were able to map the TSS and CpG islands using either the University of California Santa Cruz Genome Browser (UCSC) [35] for that species or using the UCSC liftover tool across species, providing further support that the change in XCI status was not due to differences in TSS usage.

**Figure 5:**
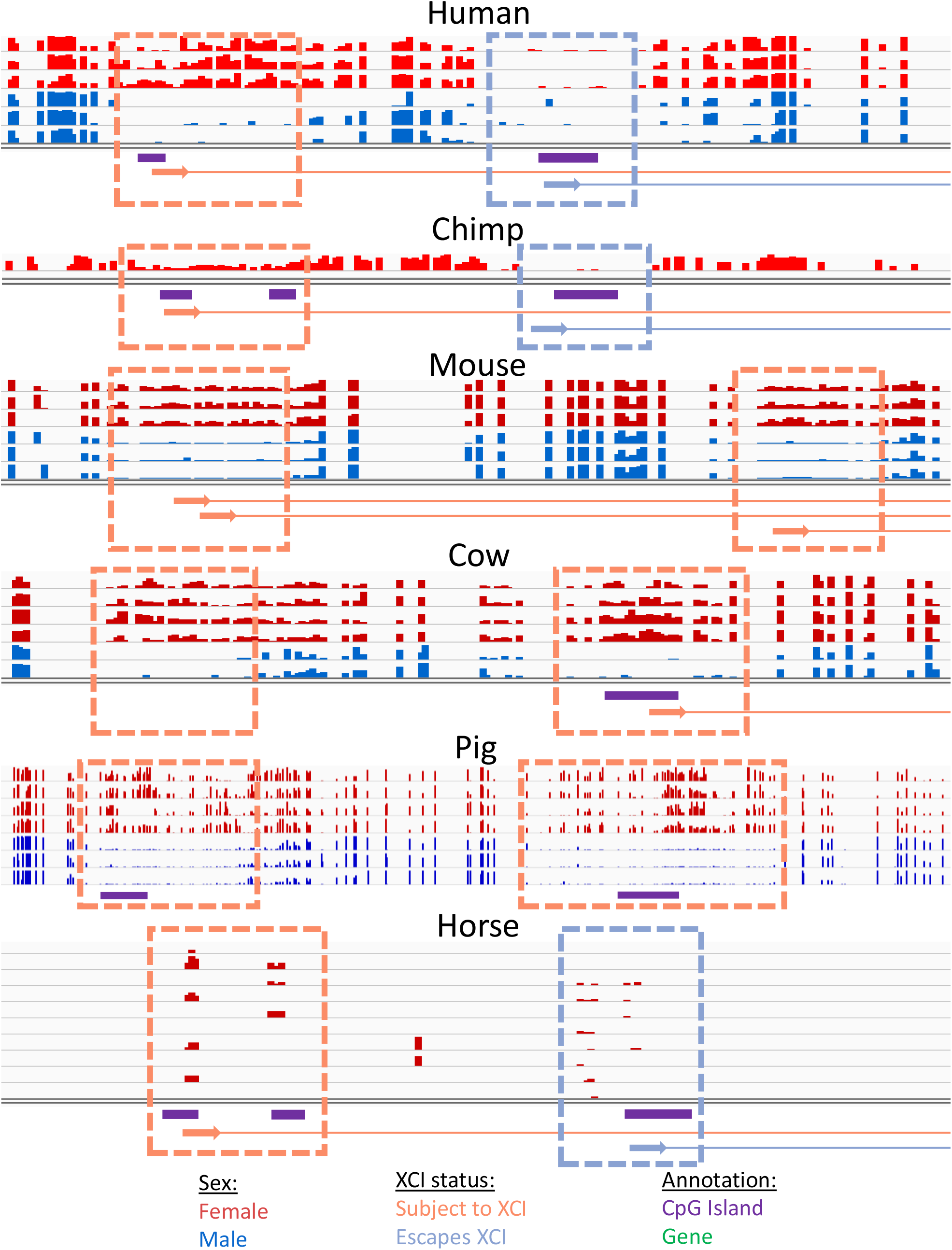
DNAme across the variably escaping gene *UBA1*. *UBA1* is featured as it has multiple different TSSs with CpG islands that have different XCI statuses. Male methylation is shown in blue and female in red. Annotated CpG islands are shown under the methylation data in orange. Genes are shown in green with arrows at the TSS pointing in the direction of transcription. All of the methylation data shown, except for horse is from WGBS. Horse used RRBS data, which is why the data is so sparse.

Looking at the position of genes escaping XCI along the human X chromosome, we saw that most genes escaping XCI clustered into domains on the short arm of the X chromosome, similar to what has been described previously [14]. Ten of the 23 transitions between clusters of genes escaping or variably escaping from XCI and genes subject to XCI fell near topologically associated domain (TAD) boundaries in human [36], again similar to what has been seen previously [37]. These clusters of genes escaping from XCI often matched across species. Genes discordant in more than one species were also often clustered, while the genes discordant in only one species were generally scattered by themselves. Some of the genes within discordant clusters were not featured in Figure 3 as they were missing data in some species. Only two of the strongly discordant genes featured in Figure 3 are located on the long arm of the X chromosome and they did not form a cluster.

We investigated these domains of changing XCI status further by examining whether the discordant species had altered the chromosomal arrangement of these genes. For the primate-specific region of genes escaping XCI spanning the genes *TCEANC* to *GEMIN8*, most species had the same gene order, orientation and flanking genes as observed for human (**Figure S6**), although some small changes were observed in gorilla, mouse, cow and sheep. In human and mouse, the two species with Hi-C data, there is a TAD spanning from *EGFL6* (which neighbors *TCEANC*) to *GEMIN8*, which may coordinate the regulation of this region, although if regulated as a domain, *EGFL6* would be expected to also escape XCI in primates. There was no data here giving an XCI status for *EGFL6*, but a previous study had seen it as subject to XCI in human [38]. Gorilla was the only primate that did not demonstrate escape from XCI across this domain, with only the gene *GEMIN8* escaping XCI. A small insertion was present in gorilla, but it was outside of the TAD which cast doubt about whether it could be the cause of this discordance from the other primates. None of the structural differences in this region were conserved across species with concordant XCI status; thus, we found no detectable genomic correlate underpinning the change in XCI status. Similar results were found for the other discordant regions.

These genes that transition their inactivation status across species provided a dataset to interrogate for factors underlying establishment of silencing or escape from silencing. We considered various factors pertaining to CpG islands in addition to enrichment of various classes of DNA repeats. No differences were seen in CpG island size, nor CpG and GC content between species with discordant XCI status at specific genes. Differences in islands between all genes escaping from versus subject to XCI per species were seen in some species, but no characteristic was seen to be significant after multiple testing correction or in more than one species.

Different classes of repeats were tested for correlation with genes escaping from versus subject to XCI in human, chimp, mouse, cow, sheep, pig and horse. There were significantly more LINE repeats within 15kb upstream of genes subject to XCI than for genes escaping from XCI in chimp, mouse, sheep and horse (**Figure 6A**, t-test, corrected *p*-values<0.01). Other repeat classes found enriched across multiple species include LTR, DNA and snRNA repeats, which were enriched at genes escaping XCI in 3 species (**Figure S7**). SINE repeats, which have previously been seen enriched at genes escaping from XCI [39], were only found significant in horse, which unexpectedly had more SINE repeats near genes subject to XCI than at genes escaping from XCI. Human still had more SINE repeats near genes escaping XCI than subject to XCI on average, but this difference failed to reach significance in this study.

**Figure 6:**
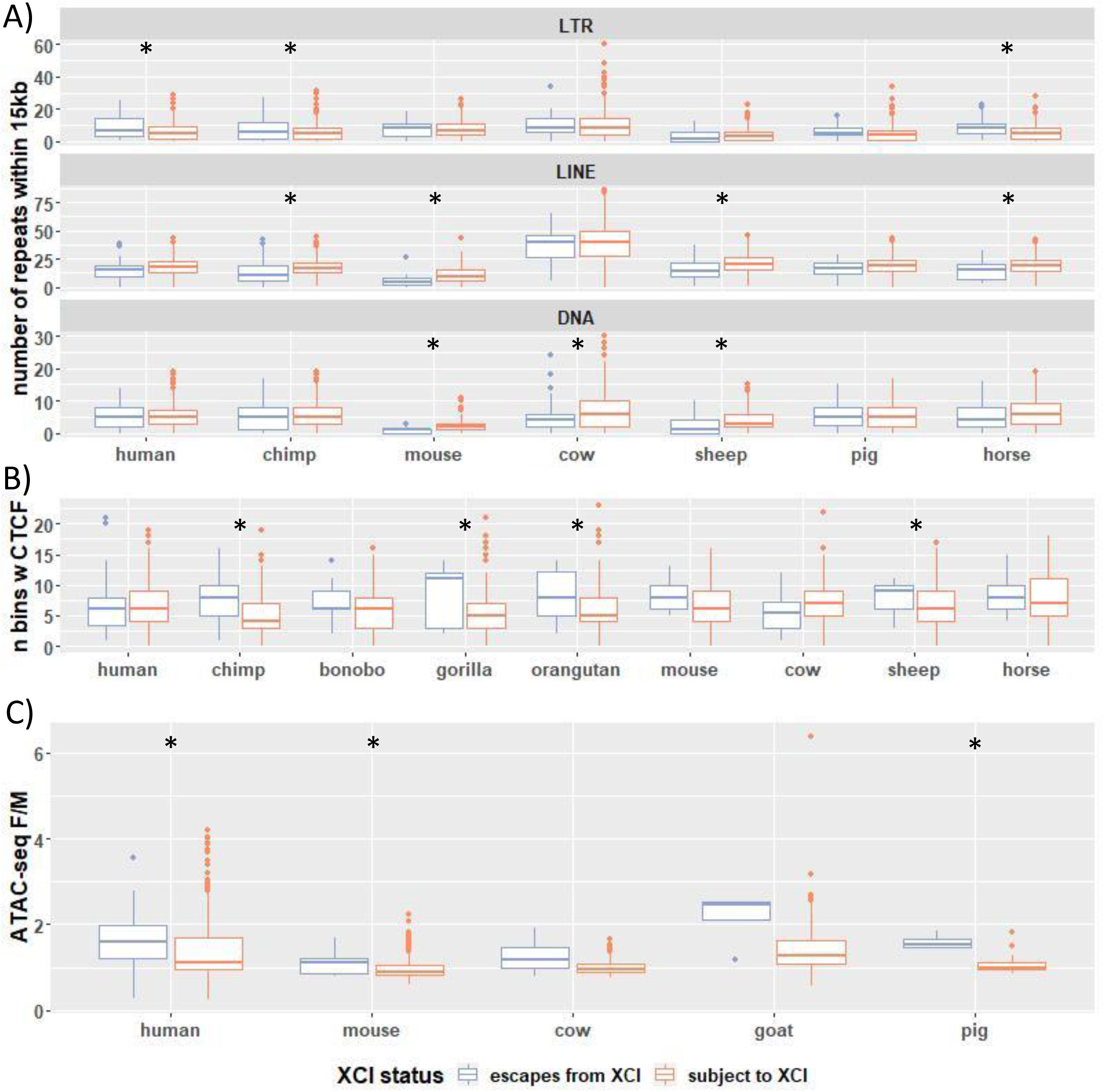
Enrichment of elements which may be related to XCI status. A) The number of repetitive elements of each class within 15kb of each gene, sorted by XCI status. See Figure S7 for the repeat classes not shown here. B) CTCF binding in overlapping 200 bp bins was predicted using a DanQ model [40]. The Y axis shows the number of bins with >50% predicted probability of having CTCF binding within 4kb of each gene. C) Female/male ATAC-seq signal averaged across samples within 250bp of each TSS. F/M is female over male. Species with a **\*** have significant differences between genes escaping XCI and those subject to XCI at adjusted p-value<0.01.

We compared CTCF binding signal between genes found escaping vs subject to XCI across species. For this, we predicted the probability of CTCF binding across species by using a DanQ model [40] trained on human CTCF ChIP data from ENCODE [41] and validated on mouse (**Figure S8**). There were significant differences in the amount of CTCF binding signal within 4kb of genes escaping vs subject to XCI in chimp, bonobo, gorilla, and horse but not in human, gorilla, mouse, cow, sheep, goat or pig (**Figure 6B**). All of the species with significant differences had more CTCF binding signal near genes escaping XCI. We also examined whether there were significant regions in the *TCEANC* to *GEMIN8* cluster of discordant genes which correlated with a change in XCI status across species but did not find any differences consistent across species (**Table S5**).

ATAC-seq is an assay for accessible chromatin [42]. Comparing ATAC-seq signal 250bp up and downstream of TSSs across species revealed significant differences in the mean female/male ratio across genes that were escaping vs subject to XCI in human, mouse and pig but not in cow or goat (**Figure 6C**). ATAC-seq signal had a higher female/male ratio in genes escaping XCI than genes subject to XCI, as seen previously in human [43], and the same trend existed in species where the differences failed to reach significance. In the species with significant differences in ATAC-seq signal with XCI status, we did not see all tissues showing significant differences (**Figure S9**). The differences were significant in the only tissue examined in human, two of the three examined in pig, and one out of ten examined in mouse.

Across all species examined, mouse genes appeared uniquely well-silenced. We clustered all species based on their XCI status calls (**Figure S10**). The bovids (cow, sheep and goat) as a group clustered together, although mouse clusters with them for an unknown reason. Dog has very sparse data which may explain it clustering as an outlier, but we are unsure of the reason why pig clustered with dog instead of with the more closely related bovids. We observed clear separation of the primates from most other species due to the large number of primate-specific escape genes.

## Discussion

Escape from XCI is an important contributor to sex differences in expression and has even been argued to underlie a male predisposition to cancer [17,28]. In addition, genes subject to XCI can also have unique effects on phenotype, with some mutations having phenotypic effects only when separate cell populations are expressing two different alleles [44,45]. Mutations that are deleterious at the cellular level or affect the region controlling choice of Xi can lead to skewed Xi choice, leaving the individual vulnerable to recessive mutations on the opposite X chromosome [46,47]. Knowing the XCI status of genes is also important for estimating the effect of an X-linked allele in genome- or epigenome-wide association studies [48,49] and is important for genetic selection of X-linked genes in agriculture [29].

To validate our use of DNAme to call XCI status, we compared expression-based calls with DNAme in human and mouse. The human Xi/Xa expression-based calls had 83% agreement with previous calls, with the discrepancies largely in genes variably escaping from XCI [15]. As cancer samples were used to allow Xi/Xa analysis, some epigenetic dysregulation may have occurred [20]. Our study was further limited by the need for heterozygous polymorphisms, thus with only 8 samples, any mis-regulation may not have been noticeable, or led to false or missed calls of variable escape from XCI. Our human DNAme calls were 94% (WGBS) and 91% (450k array) concordant with previous XCI calls, and the two datasets analyzed here gave calls that were 97% concordant with each other. Of the few XCI status calls that were inconsistent with previous studies, 80% were in genes called as variably escaping from XCI, and are likely due to differences in the population or tissues sampled. While our mouse Xi/Xa expression-based calls had a median 90% concordancy across datasets, we only identified 60-86% of previously identified mouse escape genes, likely due to differences in thresholds between studies. There were no discordancies between our mouse DNAme calls and previous mouse studies; however the genes discordant between our Xi/Xa expression calls and previous mouse studies were not informative in our DNAme calls due to lack of CpG islands. Comparing our mouse DNAme calls to a previous study by Keown *et al*., which examined DNAme on the X chromosome in mouse brain, revealed no discordancies in genes called as escaping XCI, but there were differences in which genes were informative [26].

In this study we have made an average of 342 XCI status calls per species, for 12 different species. The proportion of genes subject to XCI differs, with most species having 80-90% of genes subject to XCI. The only species with more genes subject to XCI is mouse at 95%, and the only species with fewer was horse at 76%. Additionally, horse had elevated numbers of genes variably escaping from XCI (10), while other species only had 0-5% of genes variably escaping from XCI. A meta-analysis in human found 8% of genes variably escaping from XCI and a further 7% as varying between studies [15], while our current study identified 6% variable escape in human by expression and only 2% by DNAme. Our study is consistent with a previous study using DNAme to make XCI status calls that did not see many genes consistently variably escaping from XCI [23]. Of the genes previously predicted to variably escape from XCI [15], 69% had no data in this study due to lack of a CpG island and another 10% were hypermethylated in males or females and therefore XCI status could not be determined.

Our DNAme analysis found that human genes subject to XCI have promoter CpG DNAme between 38% (in WGBS) and 41% (in 450k analysis) which agrees with a previous analysis using the 450k DNAme array which showed genes subject to XCI having an average DNAme around 40% [23] (**Table 1**). Mouse had a lower 27% DNAme average for genes subject to XCI; other mouse studies have not examined genes which are subject to XCI. Other species had DNAme averages in a range between human and mouse, but most were closer to human than mouse. Our DNAme thresholds to call genes as escaping from or subject to XCI were consistent across human and mouse WGBS but as our data was from different studies using different techniques on different tissues in different species there may be variation unaccounted for with our thresholds. However, WGBS and 450k array-based XCI status calls were consistent in both human and chimp and, with a few notable exceptions, genes had concordant XCI status calls across species. Past studies of XCI status calls using DNAme in human did not see many differences in DNAme-based XCI status across tissues [23], so different tissues analyzed may not cause many discordancies. For the primate and dog samples which used the human 450k methylation array, only probes which mapped consistently between the species were kept by the source publications [50,51], and so these species may be enriched for genes with a conserved XCI status. Utilizing datasets from different studies confounds the species differences with other experimental differences including sample size as well as inclusion of male samples. The lack of male samples in some species prohibited us from filtering out genes that are methylated on the Xa and therefore would never be seen to escape XCI by DNAme.

We compared DNA repeats and CpG island characteristics with XCI status within and across species and found none varied significantly across species per discordant gene, few varied between XCI statuses within a species and none varied between XCI statuses in all species. The most commonly enriched repeat, LINE elements, were seen enriched in genes subject to XCI within 15kb of their CpG island, similar to results seen in Cotton *et al*. (2014) [39]. The authors also saw enrichment of SINE elements at genes escaping from XCI. SINE elements were found more commonly at genes escaping XCI in human in this study, but not in significant numbers; horse, which was significant for SINE elements, had them enriched at genes subject to XCI. LTR and DNA repeats have been previously reported as contributing to a sequence-based XCI status classifier, with the LTR MLT1K being found almost exclusively within 100kb upstream of genes subject to XCI while the DNA repeat MER33 is mostly found within 100kb upstream of genes escaping XCI [52]. Here, we found the opposite phenomena, with LTRs enriched near genes escaping XCI while DNA repeats were enriched near genes subject to XCI. Neither MER33 nor MLT1K were enriched within 15kb or 100kb of CpG islands with one XCI status vs the other. We also found a novel enrichment of snRNA repeats at genes subject to XCI in multiple species, however snRNA repeats were rare with a maximum of four repeats within 15kb and often no repeats were seen in the window, while the other significant repeat classes had higher averages and ranged up to 90 repeats per 15 kb CpG island. We also predicted CTCF binding and observed that some species have more CTCF binding signal around genes escaping XCI than genes subject to XCI as has been seen previously [16,53,54]. ATAC-seq signal, which has previously been seen enriched at genes escaping XCI, was also seen enriched here, but again, only in some species [43]. A deeper bioinformatic analysis comparing our XCI status calls to features which differ across species with differing XCI status but are conserved in species with conserved XCI status might identify important regulatory features which control the XCI status of nearby genes or control XCI in general.

Our XCI status calls across species also allow us to check conservation of elements that may control XCI. A region escaping XCI in human was still able to escape from XCI when inserted at a mouse region which is normally subject to XCI, showing that the mechanisms controlling escape from XCI are conserved and functional across species [55]. We suspect that any elements found to be important in human or mouse research will be conserved across species with the same XCI status; having a variety of mammalian species with XCI status calls gives us a platform to test this hypothesis.

Many of the genes escaping from XCI have previously been seen grouped in domains [37], and here we see these domains conserved across species. Furthermore, we see that many of the genes that change XCI status across species are clustered into domains and many of these domains coincide with TADs in human. These domains suggest escape from XCI may be regulated at a domain level; however, we also see some genes being regulated individually and even separate TSSs for the same gene can have opposite XCI statuses. Individual escape genes are often discordant in a few species. Coincidence of changes in XCI status with loss of Y homology emphasizes the importance of dosage for determining genes whose escape from XCI is vital to survival. Generally, the TSS is seen to be conserved, even when a gene changes XCI status. Previous studies have suggested that CTCF and YY1 may be enriched near genes escaping from XCI [16,53,54]. CTCF has also been seen enriched at boundaries between domains of genes with opposite XCI statuses [56]. Repeat elements (SINE for genes escaping XCI and LINEs for genes subject to XCI) have also been seen enriched in 100kb windows around TSSs as well as windows 15kb upstream [39,52].

These XCI status calls may be improved in the future through new techniques such as single-cell RNA-seq (scRNA-seq) which can make expression-based XCI status calls without the need for samples with skewed Xi choice. Cells can be analyzed individually or their Xi choice can be identified and then all of the cells with the same Xi can be pooled. scRNA-seq has also identified variable escape at the cellular level within a tissue [17], with most genes varying based on their Xi choice and one gene (*TIMP1*) seen to vary randomly with no observed difference in Xi choice between cells with different XCI status. Current scRNA-seq datasets have a limitation of low read depth per cell, which limits the ability to examine lowly expressed genes [57]. Methods to enrich for the 3’ end of genes, such as the Chromium Next GEM Single Cell pipeline, are useful for quantifying expression per gene but further limits the number of polymorphisms available for study. As sequencing becomes cheaper and scRNA-seq technology continues to develop, scRNA-seq may become the new gold standard for making XCI status calls.

Non-CpG DNAme may allow us to use DNAme to examine XCI status in genes without CpG islands, as this mark is seen enriched in the gene body of transcribed genes [25]. Brain and pluripotent cells have the most abundant non-CpG DNAme, with other tissues having less than 1% non-CpG DNAme [58]. A study across multiple tissues in human found 18% of genes (109 of 612) had female-specific non-CpG DNAme in at least one tissue, but of these 66% (72 genes) were only significant in one tissue (usually brain) [27]. Another study, in brain only, found 20% of genes escaping from XCI [25]. These numbers are higher than other reports of escape, likely due to many of these genes variably escaping from XCI and only escaping from XCI in brain.

Improved gene and genome annotations in some of the less well-studied species would enhance our XCI status calls across species. Many of the species examined here had their gene annotations generated bioinformatically using CESAR [59] mapping of human genes instead of being annotated with mRNA from that species. This may not have captured the correct TSS, and if transcription was no longer close to the same CpG island these XCI status calls would be invalid. With better annotations in the future, these datasets could be reprocessed to provide more up-to-date XCI status calls with improved confidence.

As mouse has considerably fewer genes escaping from XCI than other species, there may be a better species to use as a model for research related to which genes escape from XCI. Unfortunately, none of the species other than mouse examined here are small or make affordable model systems. Rabbit, for which there was no DNAme data available, has been shown to be more similar to human than mouse in aspects of XCI and may be a good species for further examination [1].

## Conclusions

Our study has created reference XCI status calls for 12 species, so that labs working with diverse mammalian species will have improved understanding of how their genes of interest are expressed in their species of interest. We have again confirmed that mouse has substantially fewer genes escaping from XCI than human, and shown that other mammals are more similar to human in this regard. Additionally, we have shown conservation of XCI status across the majority of X-linked genes and highlighted some genes of interest which are discordant across species. Interestingly, many of these discordant genes occur in domains of similarly regulated genes. In the future, we hope to use these XCI status calls to identify elements which are controlling escape from XCI and which are conserved across species, and these discordant genes are ideal candidate regions to investigate.

## Methods

### Xi/Xa expression-based XCI status calls

Human whole genome seq and RNA-seq data was obtained for 11 samples, from the Center for Epigenome Mapping Technologies. This data is from cancer samples, and because cancer has a clonal origin, we anticipated they would show skewing of XCI. Eight of the samples had skewed Xi choice, as could be seen by the majority of genes having an Xi/Xa ratio below 0.1. These samples were from brain, blood, breast and thyroid, however neither of the brain samples had fully skewed Xi choice and could be used in this analysis. Mouse RNA-seq data was obtained from two studies using crosses between two distantly related mouse strains, one of which used an *Xist* knockout to skew Xi selection [16] and another which used fluorescent markers expressed on each X chromosome to separate cells by Xi choice [21]. These mouse datasets have previously been used to find genes escaping XCI, but most mouse studies do not call genes which are subject to XCI, so they were reanalyzed here.

The different species were processed differently due to different starting file types. The human data was pre-aligned, starting as DNA VCF files and RNA bam files. The DNA VCF files were indexed and then filtered to only heterozygous SNPs in exons using the bcftools view tool [60]. A BCF file was made for the expression data using samtools mpileup with the -t DP,AD options, followed by bcftools filter to filter for depth 30 or higher [61]. The RNA BCF file was then indexed and then bcftools call used to find indels and bcftools view used to filter for quality 30+ calls. In mouse, the data was available as fastq files and were aligned using the MEA pipeline [62]. The resulting unnormalized big wig files were then quantified at known polymorphisms to determine the number of reads on the Xi and Xa.

The levels of each allele in the RNA were then extracted using R and compared at all the heterozygous sites found in the DNA analysis [63]. The ratio between alleles was used for graphing and the error rate determined using a binomial model with an α of 0.05 [16]. Genes were assigned XCI status calls per SNP, with a ratio of 0.1 being used as a threshold between genes escaping and subject to XCI and not giving an XCI status for genes who cross this threshold with their error rates.

SNPs were mapped to splice variants which include the SNP and the closest TSS of these was used to connect DNAme and Xi/Xa expression for figures 1, S1 and S2.

### DNAme based XCI status calls

GEO was searched for all WGBS, RRBS or 450k array data that was in eutherian mammals other than mouse and human. Human data was downloaded from the International Human Epigenomics Consortium (IHEC) [64], while a single mouse dataset with a high number of samples was downloaded [65]. Data was downloaded for *Homo sapiens* (human), *Pan troglodytes* (chimp), *Pan paniscus* (bonobo), *Gorilla gorilla* and *Gorilla beringei* (gorilla), *Pongo pygmaeus* and *Pongo abelii* (orangutan), *Mus musculus* (mouse), *Bos Taurus* (cow), *Ovis aries* (sheep), *Capra aegagrus hircus* (goat), *Sus scrofa* (pig), *Equus ferus caballus* (horse) and *Canis familiaris* (dog). When processed bigwig files were available they were chosen over processing from raw data. Relevant genomes were downloaded from UCSC (**Table S3**) and raw reads were aligned to them using BISMARK [66]. BISMARK methylation extractor was used to get bedGraph files and then UCSC tools bedGraphToBigWig tool used to make bigwig files. Gene and CpG island maps were downloaded from UCSC, and the UCSC tools bigWigAverageOverBed tool was used to quantify the mean methylation level across CpG islands. R was then used to annotate CpG islands within 2kb of a gene’s TSS as belonging to that gene and XCI status calls were made, with islands with a mean DNAme below 10% being called as escaping XCI and islands with between 15 and 60% DNAme being called as subject to XCI. Islands for which over half of males had 15% DNAme or higher were discarded as having male hypermethylation and being uninformative. The mean DNAme across each sex was also calculated and compared per CpG island.

For datasets generated on the human 450k DNAme array, data was downloaded and filtered for promoter associated probes. The mean DNAme of probes sharing an annotated CpG island were matched to their annotated genes and this was used for making XCI status calls as above.

### Clustering

XCI calls per species were transformed into numeric values, with escape as 0, variable escape as 0.5 and subject to XCI as 1. The daisy function from the cluster package in R was used to compute distance and then hclust with the gower metric and complete method were used to perform the clustering. The phylogenetic tree was generated using the online interactive Tree of Life tool [67].

### Conservation analysis

R was used to collect and match all the XCI status calls across species. Genes were matched based on their name, controlling only for capitalization changes across species. Genes with XCI status calls in four or more species were included in further analysis. Datasets analyzed were split into two different groups: all mammals (human, chimp, mouse, cow, pig, sheep, and goat WGBS data, with horse RRBS and dog 450k array data) and primates (human, chimp, bonobo, gorilla and orangutan 450k array data). The two separate groups allowed us to examine conservation of genes without our analyses being biased toward primate specific calls.

### Statistical tests

Statistical tests comparing enrichment of CpG island statistics and various repeat classes between genes subject to or escaping from XCI were done using R. We used a t-test with the Benjamini Hochberg method for multiple testing correction [68].

### Domain analysis

Domains were identified based on conservation calls above and examined using the UCSC browser to compare the arrangement of genes. TAD boundaries were taken from Dixon, 2012 [36] and were annotated to genes if they were between it and the next gene or were within the gene body.

### ATAC-seq analysis

ATAC-seq data was downloaded, see table S3 for data sources. If bigwig files were available they were used, but if not we downloaded raw data and aligned it using HISAT2 [69]. The bamcoverage tool from the deepTools package [70] was used to generate bigwig files (normalized using RPKM) and bigWigAverageOverBed from UCSC utilities was used to determine the mean coverage in 250bp up and downstream of each TSS. Each TSS was matched to the closest CpG island within 2kb and any XCI status call from that island used for the TSS.

### CTCF predictions

CTCF binding was predicted using a strand-specific DanQ model [40]. The model was trained on human CTCF ChIP-seq data (*i*.*e*. positive sequences) and DNase I hypersensitive sites (*i*.*e*. negative sequences) from ENCODE [71]. Presence of a CTCF binding site on the forward strand was required for positive sequences. Negative sequences were required to match the distribution of %GC content of positive sequences. To evaluate the ability of the model to make CTCF binding predictions in species other than human, it was validated on mouse CTCF ChIP-seq data (also from ENCODE). We used this model to predict the probability of having a CTCF-bound region per overlapping 200bp bins in all species but dog (which had very few XCI status calls to compare to). The CTCF model, the data used to train and validate it, and the cross-species CTCF-binding predictions on the X chromosomes of the studied species have been deposited on GitHub (https://github.com/wassermanlab/CTCF/). For the purpose of quantifying CTCF binding signal per TSS, we counted the number of bins with an over 50% predicted probability of being a CTCF-bound region within 4kb of each TSS. For our analysis of the *TCEANC* to *GEMIN8* region, we counted the number of bins with over 50% probability of CTCF within each region.

## List of abbreviations

450k: Illumina Infinium Human Methylation450 BeadChip array
DNAme: DNA methylation
RRBS: reduced representation bisulfite sequencing
scRNA-seq: single cell RNA sequencing
TAD: topologically associated domain
TSS: transcription start site
UCSC: University of California Santa Cruz Genome Browser
WGBS: whole genome bisulfite sequencing
Xa: active X chromosome
XCI: X-chromosome inactivation
Xi: inactive X chromosome

## Declarations

### Ethics approval and consent to participate

The human expression data and some of the human WGBS data was generated by The Canadian Epigenetics, Epigenomics, Environment and Health Research Consortium (CEEHRC) initiative funded by the Canadian Institutes of Health Research (CIHR), Genome BC, and Genome Quebec. Ethics approval for data access was provided by the University of British Columbia Clinical Research Ethics Board (H17-01363).

### Consent for publication

Not applicable.

### Availability of data and material

See Supplemental Table S3 and references for data sources used. All sources are publicly available.

### Authors’ contributions

BPB conducted the analyses, OF provided the CTCF prediction modelling, and all authors contributed to the interpretation of data and have approved the manuscript for publication.

## Acknowledgements

We thank the other members of the Brown and Wasserman labs for helpful comments during the development of this project.

The human expression data and some of the human WGBS data was generated by The Canadian Epigenetics, Epigenomics, Environment and Health Research Consortium (CEEHRC) initiative funded by the Canadian Institutes of Health Research (CIHR), Genome BC, and Genome Quebec. Information about CEEHRC and the participating investigators and institutions can be found at http://www.cihr-irsc.gc.ca/e/43734.html. We would also like to thank the research groups which generated the other sources of data used in this analysis.

## Tables

**Table S1: All XCI status calls made in this study compared to human**. On the far right are columns tabulating XCI status calls for the different groups divided into in our analysis.

**Table S2: Individual XCI status calls per dataset**. Each sheet is a separate dataset analyzed.

**Table S3: The sources of data used in this study**. See separate sheets for different types of data used.

**Table S4: Enrichment of repeats at genes escaping from vs subject to XCI. T**he number of repeats of each class (sheet 1) and family (sheet 2) within 15kb upstream of each gene were quantified. T-tests were done to compare the enrichment of each repeat class in genes with one XCI status vs the other.

**Table S5: The number of predicted CTCF binding sites between genes in a discordant region**. A DanQ model was given overlapping 200bp bins of each genome and predicted the likelihood of it containing a CTCF binding site. The number of bins with over 80% chance of having CTCF binding were counted per region. Each region goes from either the start of a gene to its end, or from the end of one gene to the start of the next. Edges were included 5kb from the furthest gene on each side. This discordant region is the one featured in figure S6. The mean value region species for each region.

## Figures

**Figure S1: The Xi/Xa expression ratio vs promoter DNAme level in individual human samples**. Each point is a SNP with Xi/Xa expression data, matched to the most likely promoter and any CpG islands within 2kb in order to have matched DNAme values. Lines are drawn at 0.1 Xi/Xa expression and at 10, 15 and 60% DNAme as they were used as thresholds to call XCI escape status later. Points are colored based on their XCI status calls in the previous literature [15]. CEMT30, a leukemia cancer sample, was used for Figure 1. Three samples (CEMT19, CEMT23 and CEMT43) were discarded from downstream analyses, because they did not appear to show skewing of Xi choice, with many genes called as subject to XCI by DNAme and previous studies, with an XiXa expression ratio >>0.1.

**Figure S2: The Xi/Xa expression ratio vs promoter DNAme level in individual mouse samples**. Each point is a SNP with Xi/Xa expression data, matched to the most likely promoter and any CpG islands within 2kb in order to have matched DNAme values. Lines are drawn at 0.1 Xi/Xa expression and at 10, 15 and 60% DNAme as they were used as thresholds to call XCI escape status later. Points are colored based on their XCI status calls made using Xi/Xa expression. Data from 2 different studies are used: one used an *Xist* knockout to skew Xi choice and the other used differently colored fluorescent proteins expressed from each X chromosome to sort cells based on Xi choice. Data from Keown *et al*. not shown here was used for Figure 1.

**Figure S3: Male vs female DNAme across species**. The DNAme data shown was generated with 3 different methods: WGBS, RRBS and the human 450k DNAme array. Each point is a CpG island. Lines are drawn at female DNAme of 10,15 and 60 as those thresholds were used to call a gene’s XCI status and at male DNAme of 15 as genes with higher than 15% male DNAme were discarded from further analysis.

**Figure S4: A comparison of imprinted genes and genes subject to XCI**. The average DNAme level at promoter CpG islands are shown for 4 imprinted genes and 4 genes subject to XCI in human (A) and mouse (B). Genes subject to XCI have males and females separate as females are expected to be hemi-methylated while males are expected to have low methylation.

**Figure S5: Comparison of DNAme data generated using WGBS and the 450k array**. Human and chimp were the only two species which had data generated using both methods. Lines are drawn at 10,15 and 60% DNAme to show the thresholds used for calling XCI status. Another line was drawn along the diagonal to show where perfect concordance between datasets would be. The R^2^ value was calculated showing the level of concordance found between the 2 methods.

**Figure S6: Cross-species comparison of a primate-specific escape domain**. The domain spanning from *TCEANC* to *GEMIN8* and the neighboring gene on each side are shown. Genes names are colored by their XCI status in each species and the gene diagram is colored by whether the gene annotation is from mRNA in that species or from other species. All regions in all species were scaled together, with species aligned at the end of *GPM6B*. As there is a large gene-free region between *GEMIN8* and *GLRA2* this region has been condensed and the distance between the two genes noted. Dotted lines show a region that is inverted in sheep. Xpter and Xqter show the direction to the short and long arms of the chromosome respectively, note that this region and much of the X chromosome is inverted in mouse and cow [18,72,73]. Cow had inconsistencies between bosTau6 (used in our data source and this study) and bosTau9 (the latest cow genome build), with bosTau6 being used here. bosTau9 had duplication or rearrangement of *EGFL6 and TCEANC*. Gorilla and horse had small pseudogene insertions in the region, but these were only around 2kb in size and so were left out.

**Figure S7: Number of repeats within 15kb per TSS for genes subject or escaping XCI across species**. Species with a **\*** have significant differences between genes found escaping XCI and those found subject to XCI at adjusted p-value<0.01.

**Figure S8: Tests on mouse CTCF of our model trained on human CTCF**. This is a DanQ model trained on human CTCF ChIP data from ENCODE and tested on mouse data from ENCODE.

**Figure S9: Mean female/male ATAC-seq signal across samples within 250bp of TSSs, separated by tissue**. Tissues with a **\*** have significant differences between genes found escaping XCI and those found subject to XCI at adjusted p-value<0.01.

**Figure S10**: **Clustering of species by XCI status calls**. Species were clustered by their XCI status calls (A) and compared to a phylogenetic tree showing their evolutionary relations (B). For the clustering, species names are colored by the type of data used to generate the XCI status calls.

