## Supplemental Figures for "Cross-species examination of X-chromosome inactivation highlights domains of escape from silencing"

### Supplemental Tables

**Table S1: All XCI status calls made in this study compared to humans.** On the far right are columns tabulating XCI status calls for the different groups divided into in our analysis.

**Table S2: Individual XCI status calls per dataset.** Each sheet is a separate dataset analyzed.

**Table S3: The sources of data used in this study.**

**Table S4: Enrichment of repeats at genes escaping from vs subject to XCI. T**he number of repeats of each class (sheet 1) and family (sheet 2) within 15kb upstream of each gene were quantified. T-tests were done to compare the enrichment of each repeat class in genes with one XCI status vs the other.

**Table S5**: **The number of predicted CTCF binding sites between genes in a discordant region.** A DanQ model was given overlapping 200bp bins of each genome and predicted the likelihood of it containing a CTCF binding site. The number of bins with over 80% chance of having CTCF binding were counted per region (A). Each region goes from either the start of a gene to its end, or from the end of one gene to the start of the next. Edges were included 5kb from the furthest gene on each side. This discordant region is the one featured in figure S6. The mean value region species for each region (B)

### Figure S1

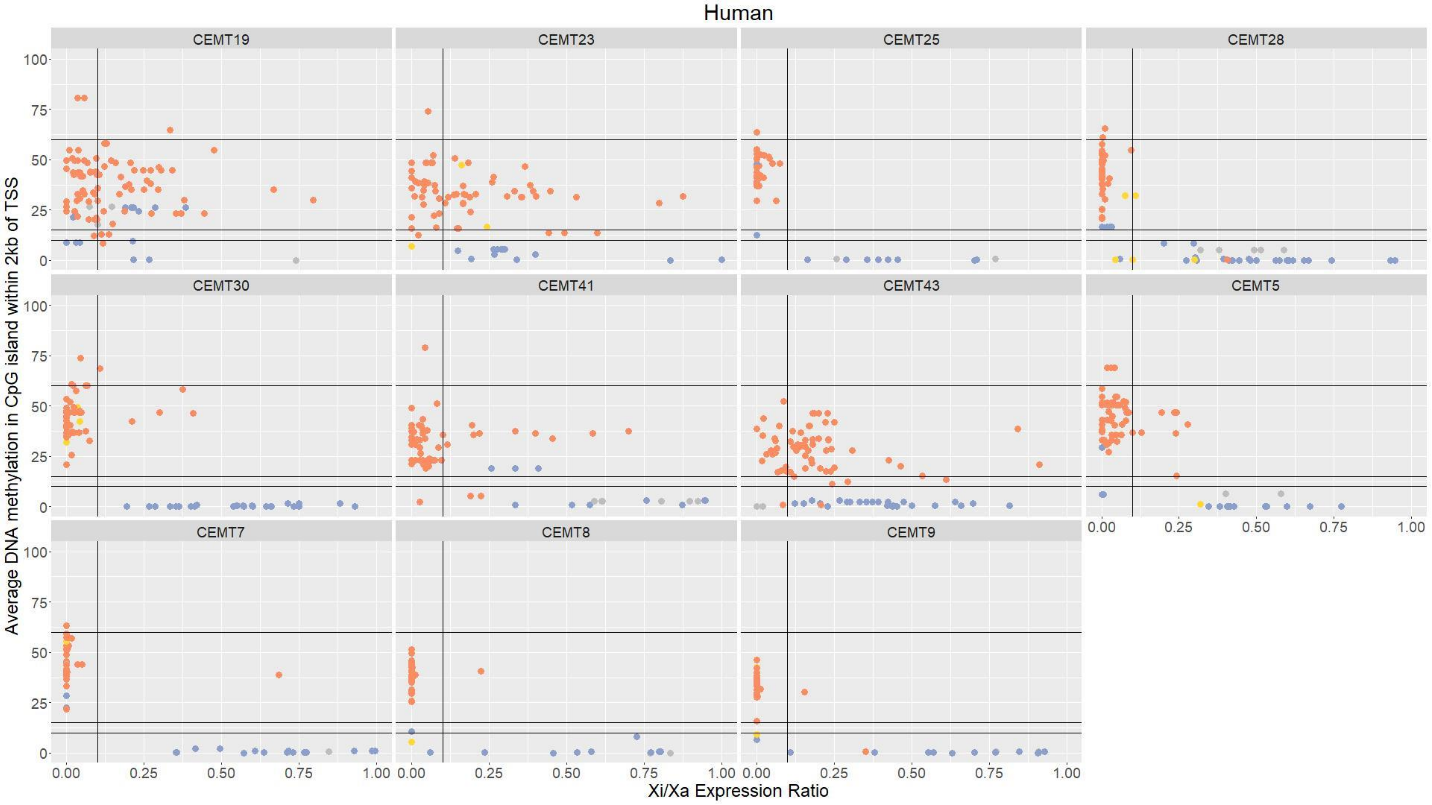


XCI status of past studies:

Escapes XCI  Subject to XCI  Variably escapes XCI

**Figure S1: The Xi/Xa expression ratio vs promoter DNAme level in individual human samples.** Each point is a SNP with Xi/Xa expression data, matched to the most likely promoter and any CpG islands within 2kb in order to have matched DNAme values. Lines are drawn at 0.1 Xi/Xa expression and at 10, 15 and 60% DNAme as they were used as thresholds to call XCI escape status later. Points are colored based on their XCI status calls in the previous literature (Balaton, 2015). CEMT30, a leukemia cancer sample, was used for Figure 1. Three samples (CEMT19, CEMT23 and CEMT43) were discarded from downstream analyses, because they did not appear to show skewing of Xi choice, with many genes called as subject to XCI by DNAme and previous studies, with an XiXa expression ratio >>0.1.

### Figure S2

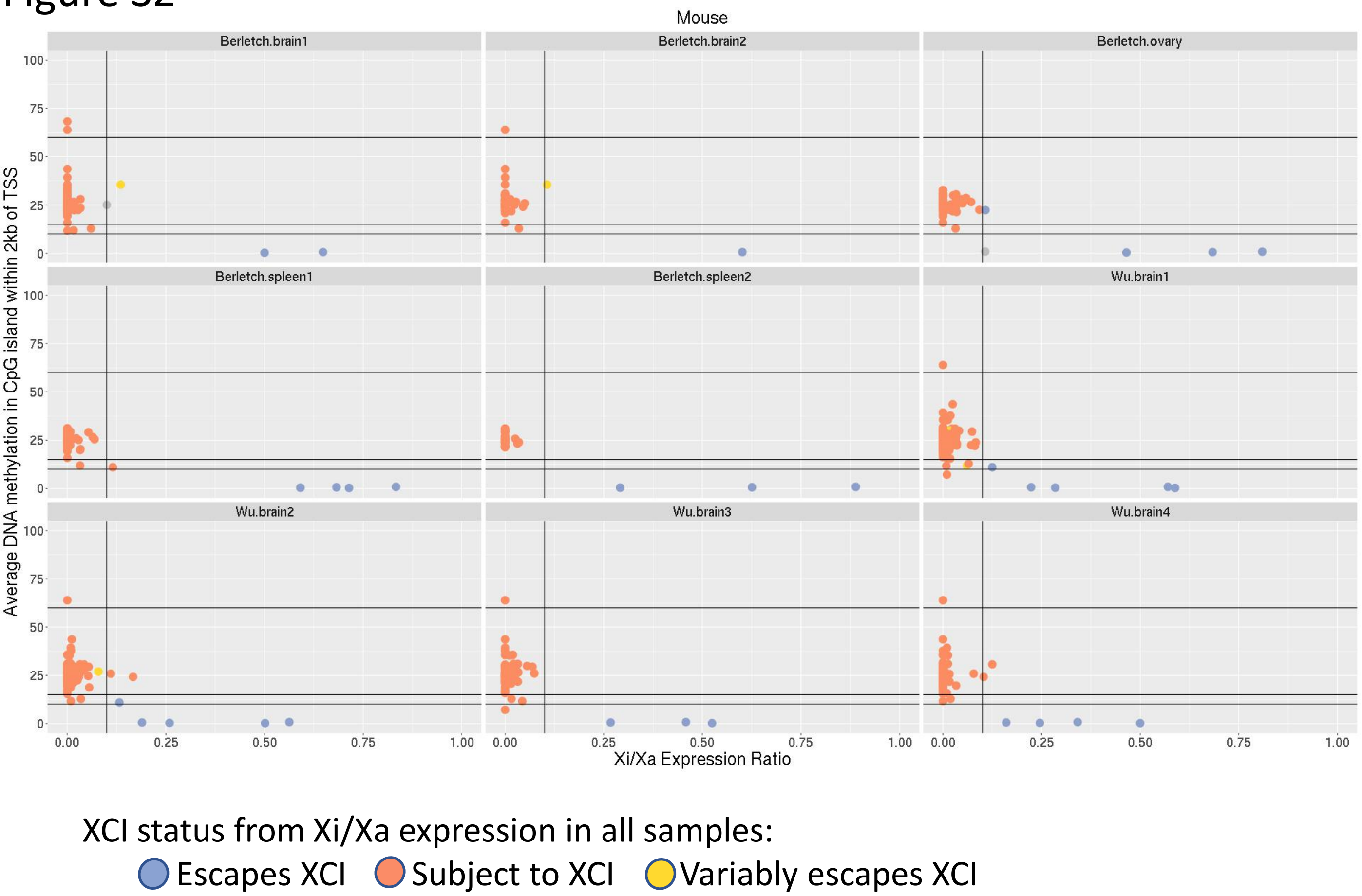


**Figure S2: The Xi/Xa expression ratio vs promoter DNAme level in individual mouse samples.** Each point is a SNP with Xi/Xa expression data, matched to the most likely promoter and any CpG islands within 2kb in order to have matched DNAme values. Lines are drawn at 0.1 Xi/Xa expression and at 10, 15 and 60% DNAme as they were used as thresholds to call XCI escape status later. Points are colored based on their XCI status calls made using Xi/Xa expression. Data from 2 different studies are used: one used an *Xist* knockout to skew Xi choice and the other used differently colored fluorescent proteins expressed from each X chromosome to sort cells based on Xi choice. Data from Keown, *et al*. not shown here was used for Figure 1.

Fig S3. **Male vs Female Methylation shows similarities across species**

Not pictured chimp (WGBS) and goat, due to lack of male data

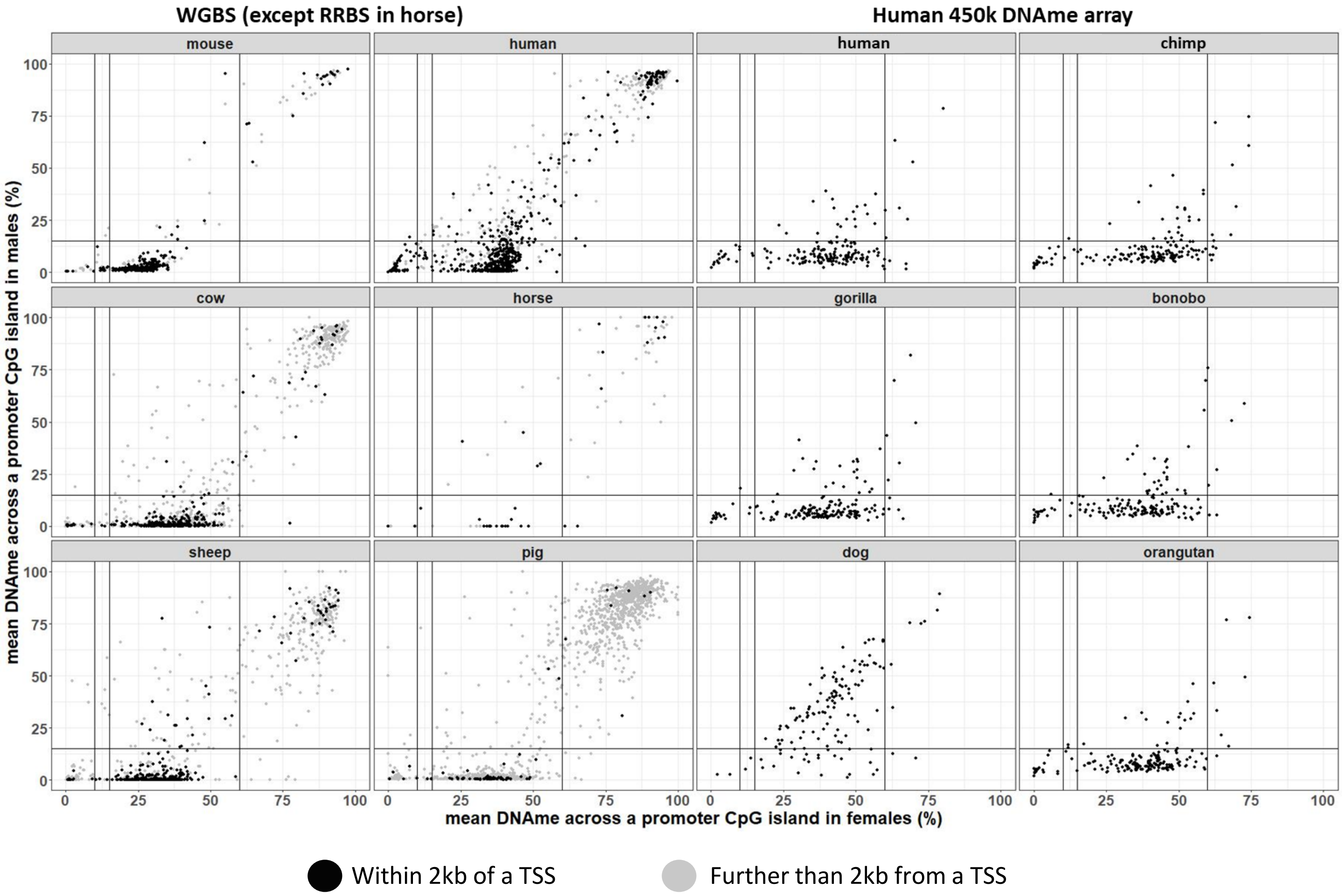


**Figure S3: Male vs female DNAme across species.** The DNAme data shown was generated with 3 different methods: WGBS, RRBS and the human 450k DNAme array. Each point is a CpG island. Lines are drawn at female DNAme of 10,15 and 60 as those thresholds were used to call a gene's XCI status and at male DNAme of 15 as genes with higher than 15% male DNAme were discarded from further analysis.

Fig S4

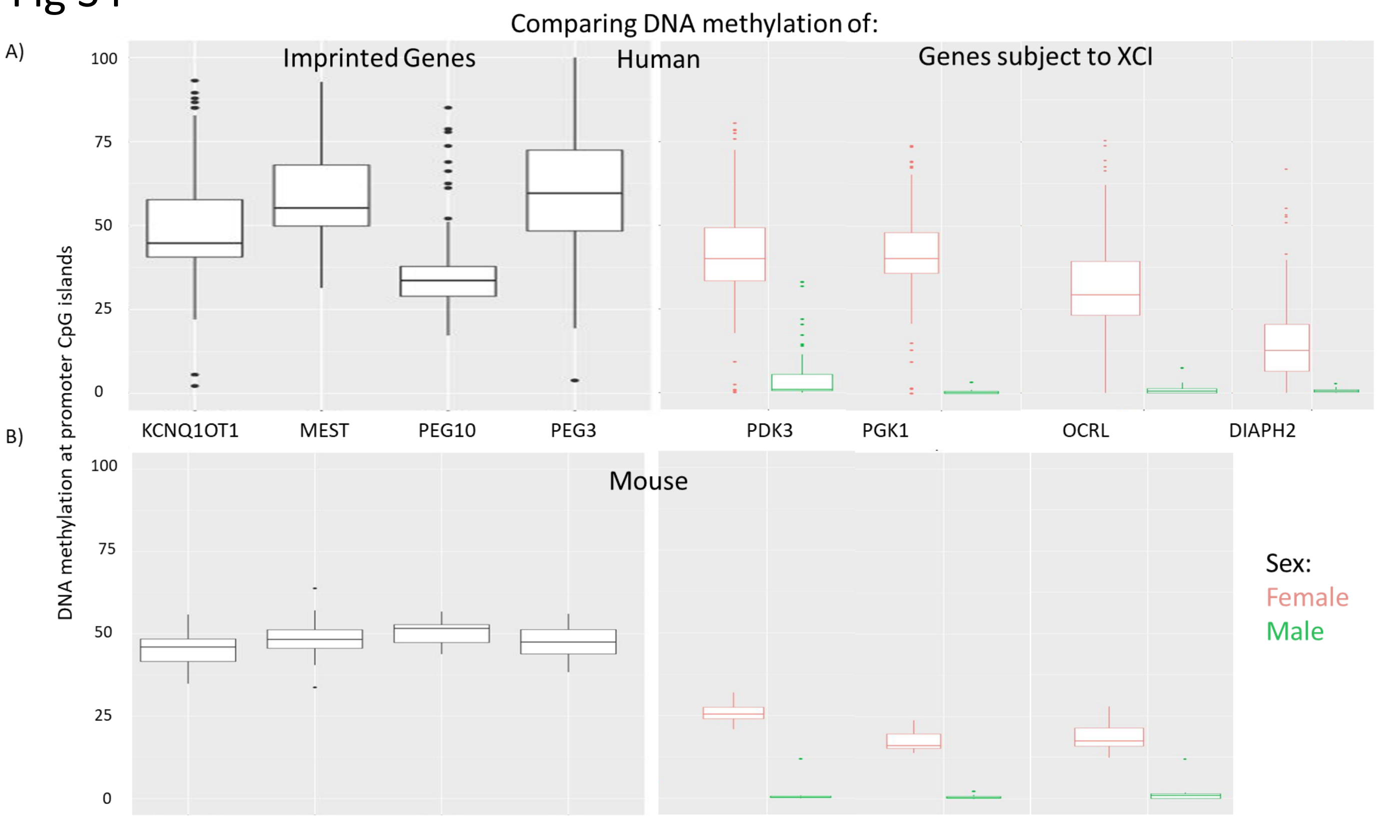


**Figure S4: A comparison of imprinted genes and genes subject to XCI.** The average DNAme level at promoter CpG islands are shown for 4 imprinted genes and 4 genes subject to XCI in humans (A) and mouse (B). Genes subject to XCI have males and females separate as females are expected to be hemi-methylated while males are expected to have low methylation.

Fig S5

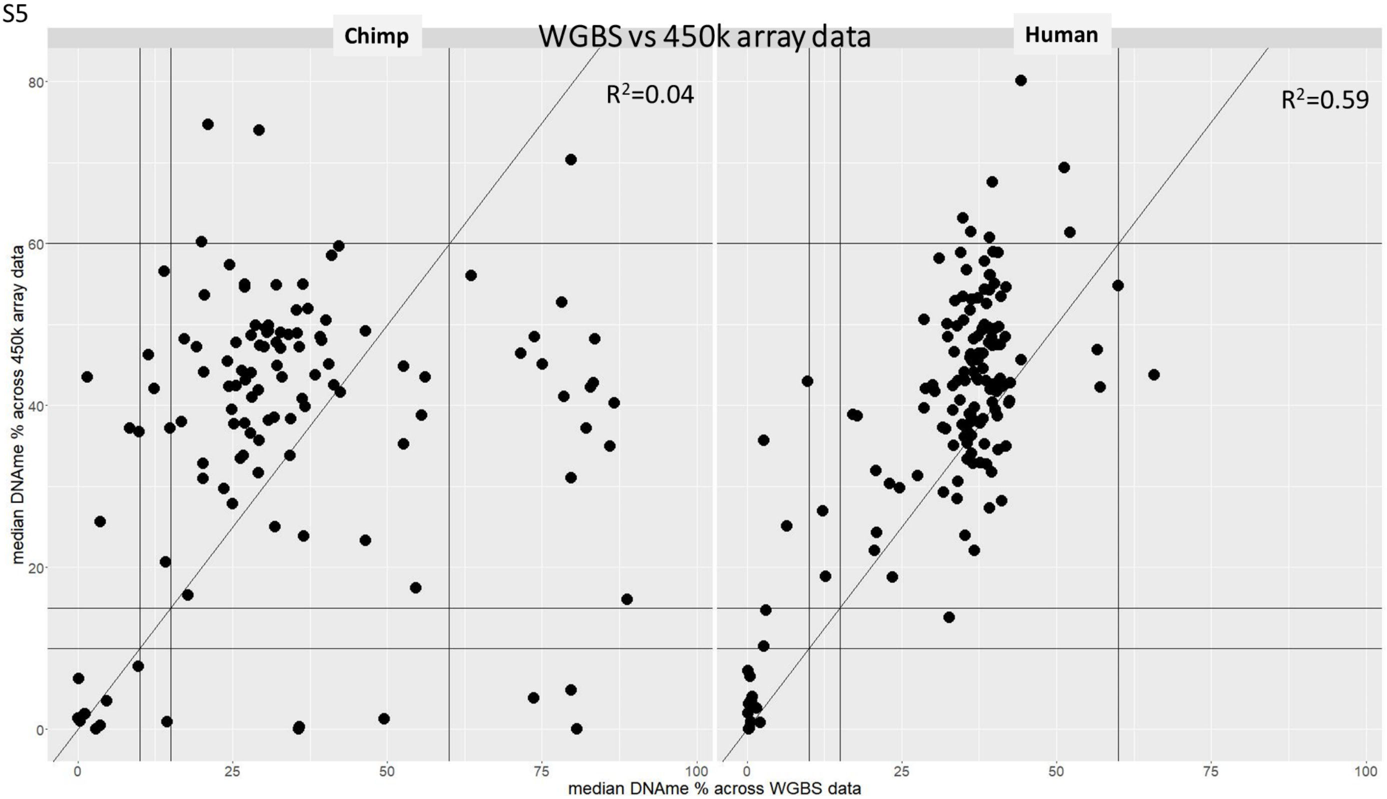


**Figure S5: Comparison of DNAme data generated using WGBS and the 450k array.** Human and chimp were the only two species which had data generated using both methods. Lines are drawn at 10,15 and 60% DNAme to show the thresholds used for calling XCI status. Another line was drawn along the diagonal to show where perfect concordance between datasets would be. The $R^2$ value was calculated showing the level of concordance found between the 2 methods.

Fig S6

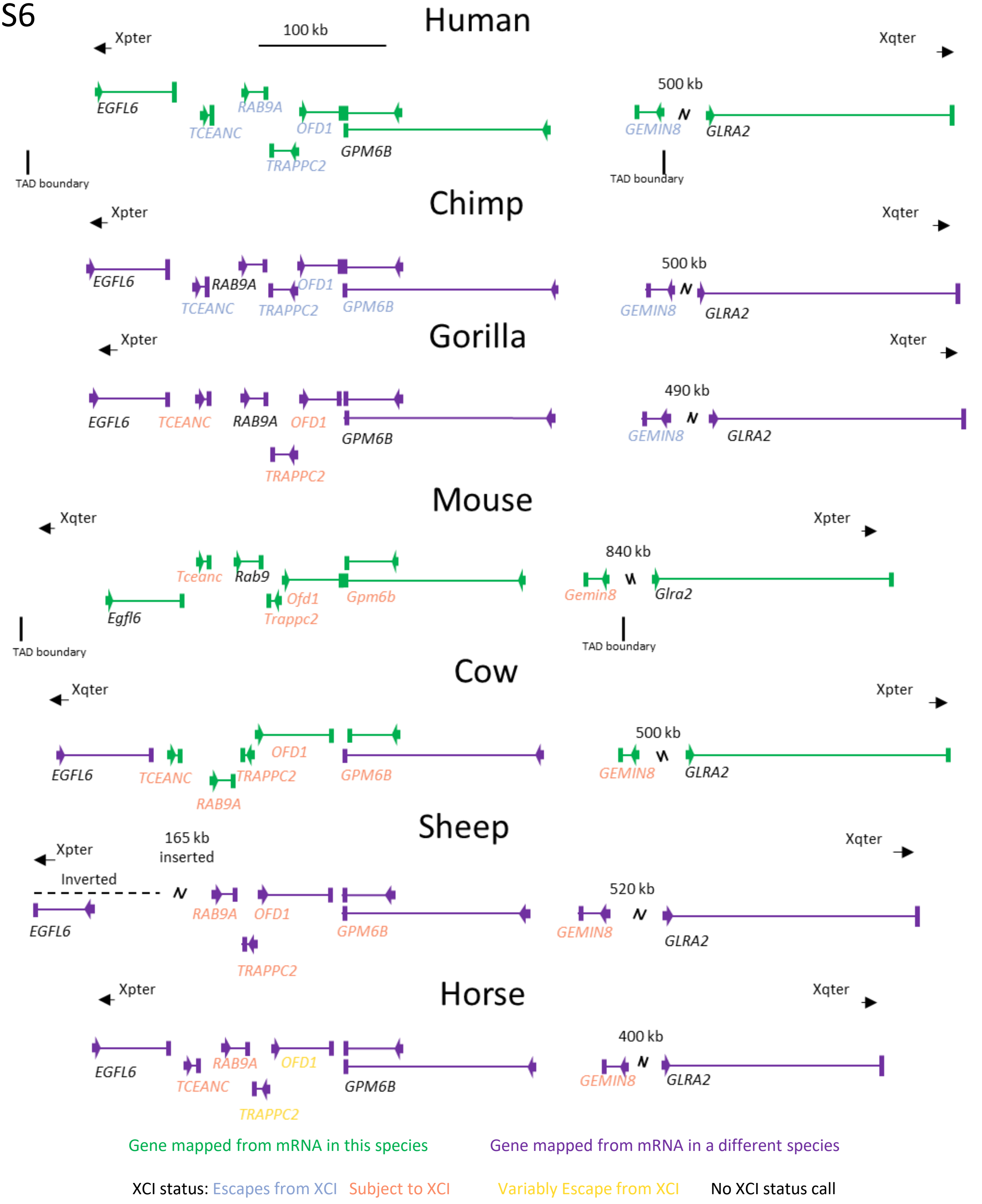


**Figure S6: Cross-species comparison of a primate-specific escape domain**. The domain spanning from *TCEANC* to *GEMIN8* and the neighboring gene on each side are shown. Genes names are colored by their XCI status in each species and the gene diagram is colored by whether the gene annotation is from mRNA in that species or from other species. All regions in all species were scaled together, with species aligned at the end of *GPM6B*. As there is a large gene-free region between *GEMIN8* and *GLRA2* this region has been condensed and the distance between the two genes noted. Dotted lines show the region that is inverted in sheep. Xpter and Xqter show the direction to the short and long arms of the chromosome respectively, note that this region and much of the X chromosome is inverted in mouse and cow [19,64,65]. Cow had inconsistencies between bosTau6 (used in our data source and this study) and bosTau9 (the latest cow genome build), with bosTau6 being used here. bosTau9 had duplication or rearrangement of *EGFL6 and TCEANC*. Gorilla and horse had small pseudo-gene insertions in the region, but these were only around 2kb in size and so were left out.

Fig S7

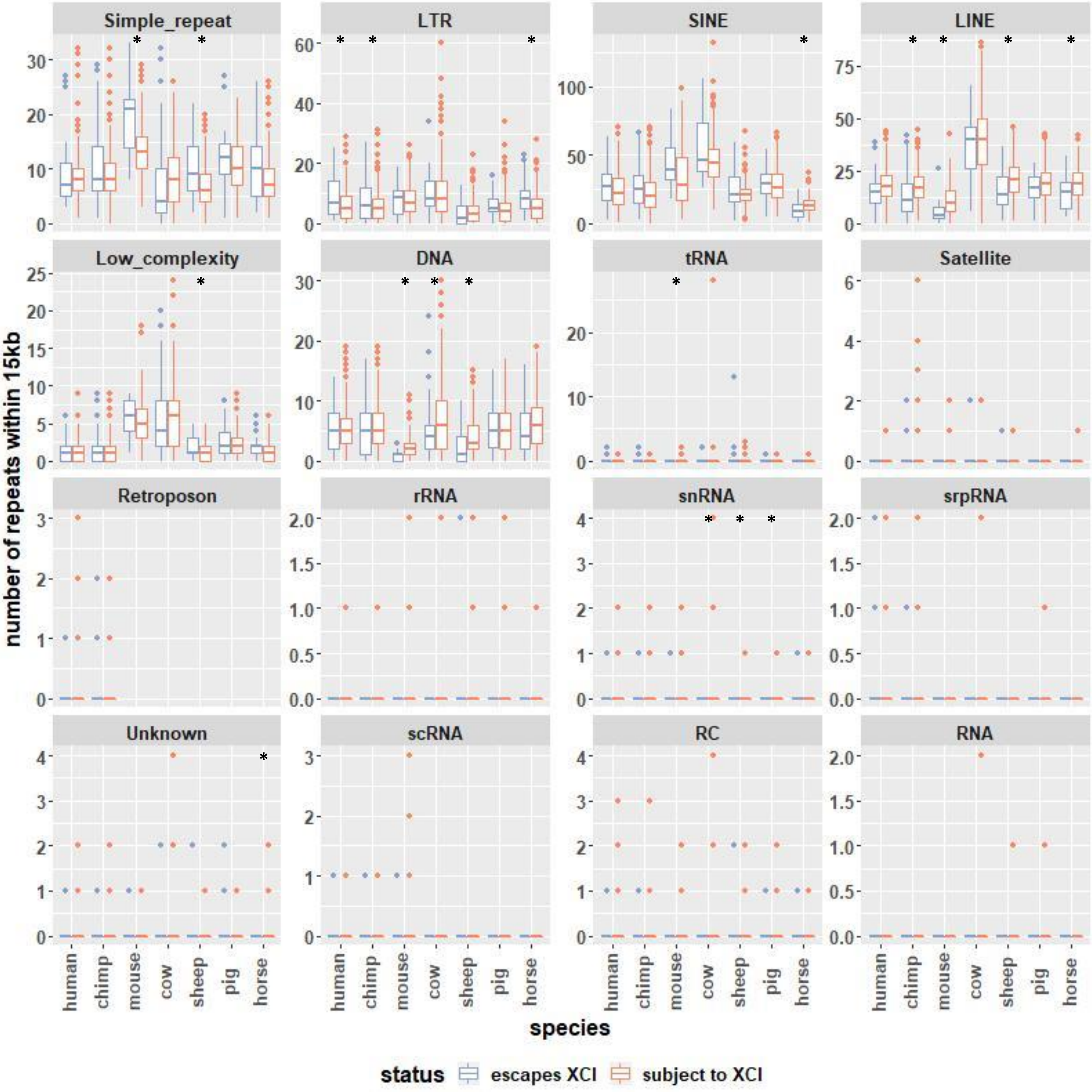


**Figure S7: Number of repeats within 15kb per TSS.** Species with a ***** have significant differences between genes found escaping XCI and those found subject to XCI at adjusted p-value<0.01.

Fig S8

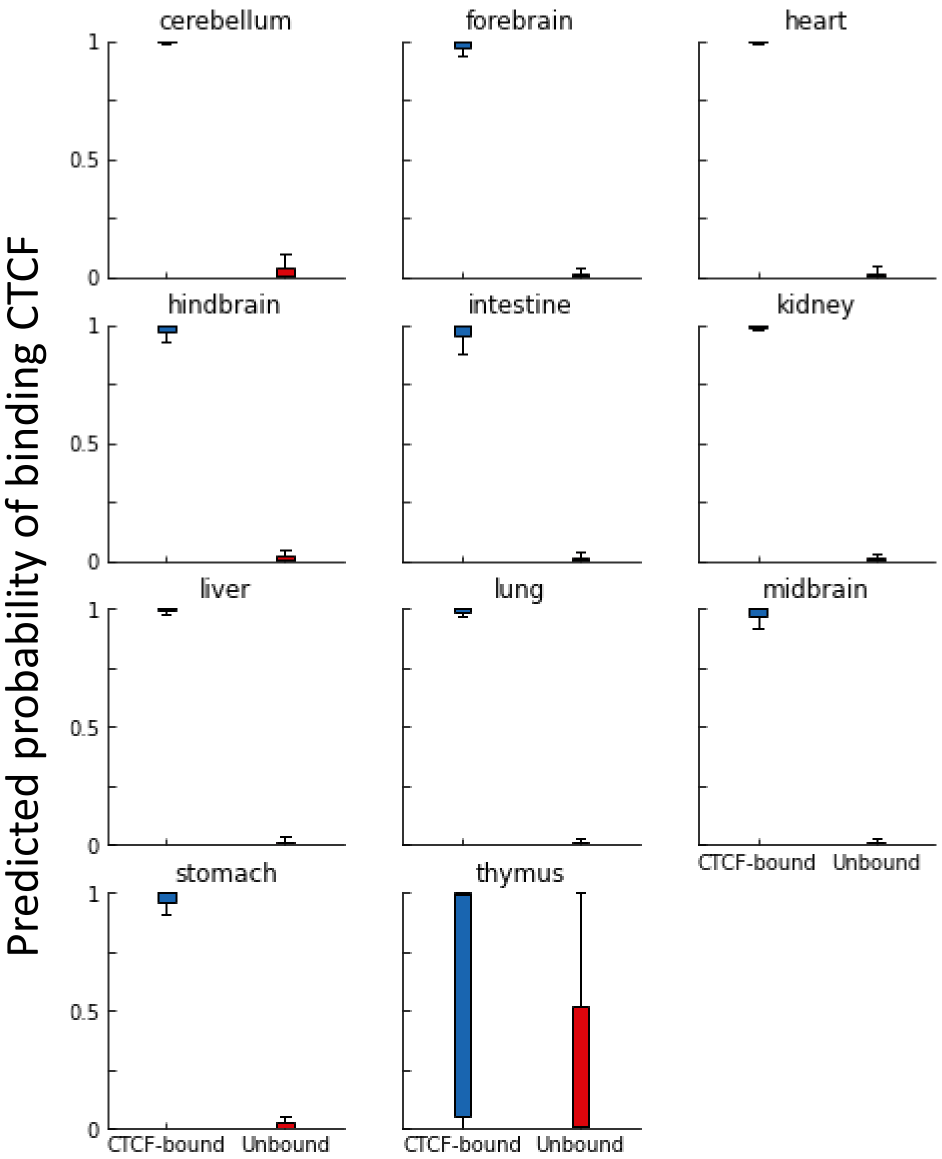


**Figure S8: Tests on mouse CTCF of our model trained on human CTCF.** This is a DanQ model trained on human CTCF ChIP data from ENCODE and tested on mouse data from ENCODE.

Fig S9

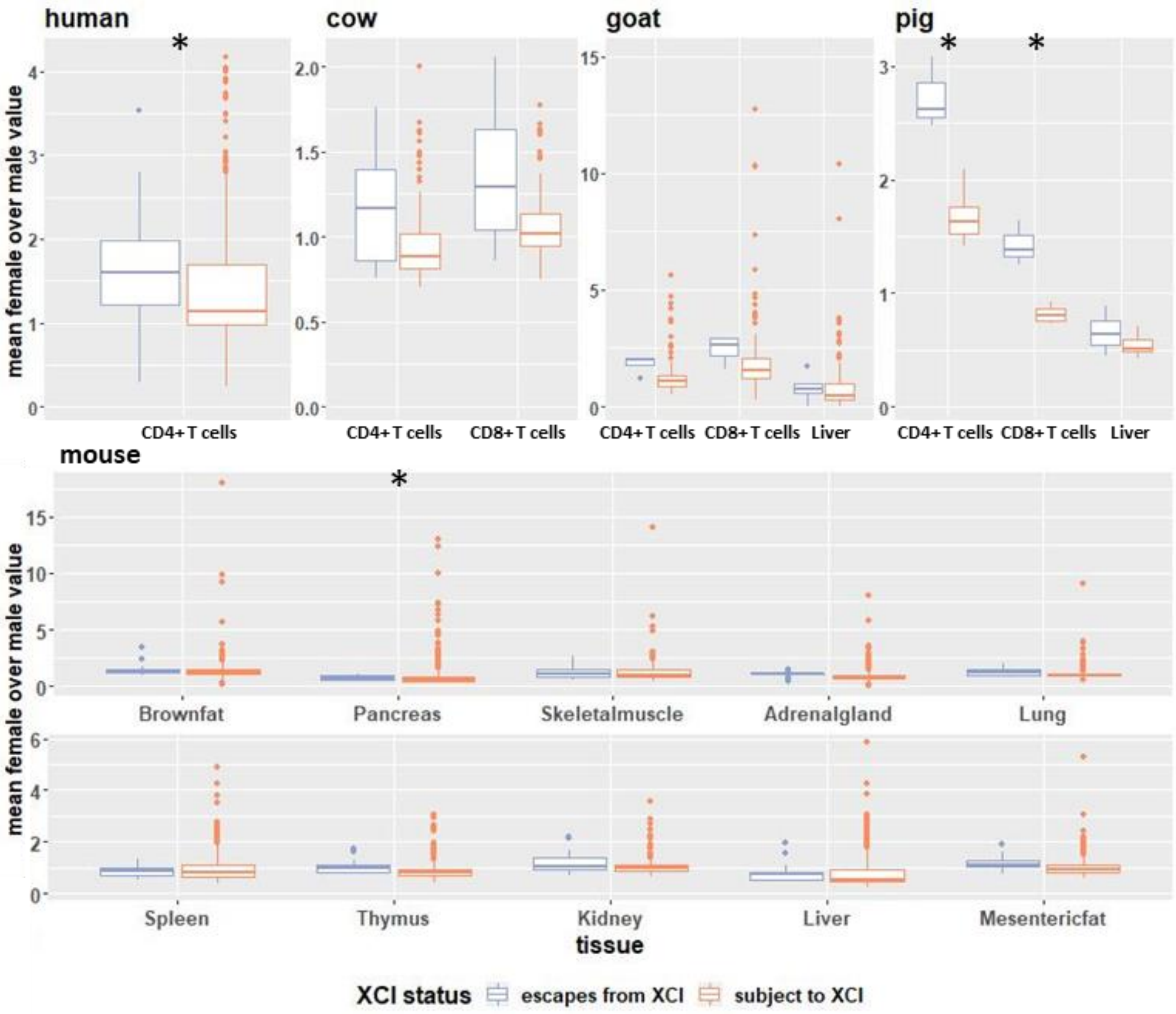


**Figure S9**: **Mean female/male ATAC-seq signal across samples within 250bp of TSSs, separated by tissue.** Tissues with a * have significant differences between genes found escaping XCI and those found subject to XCI at adjusted p-value<0.01.

Fig S10

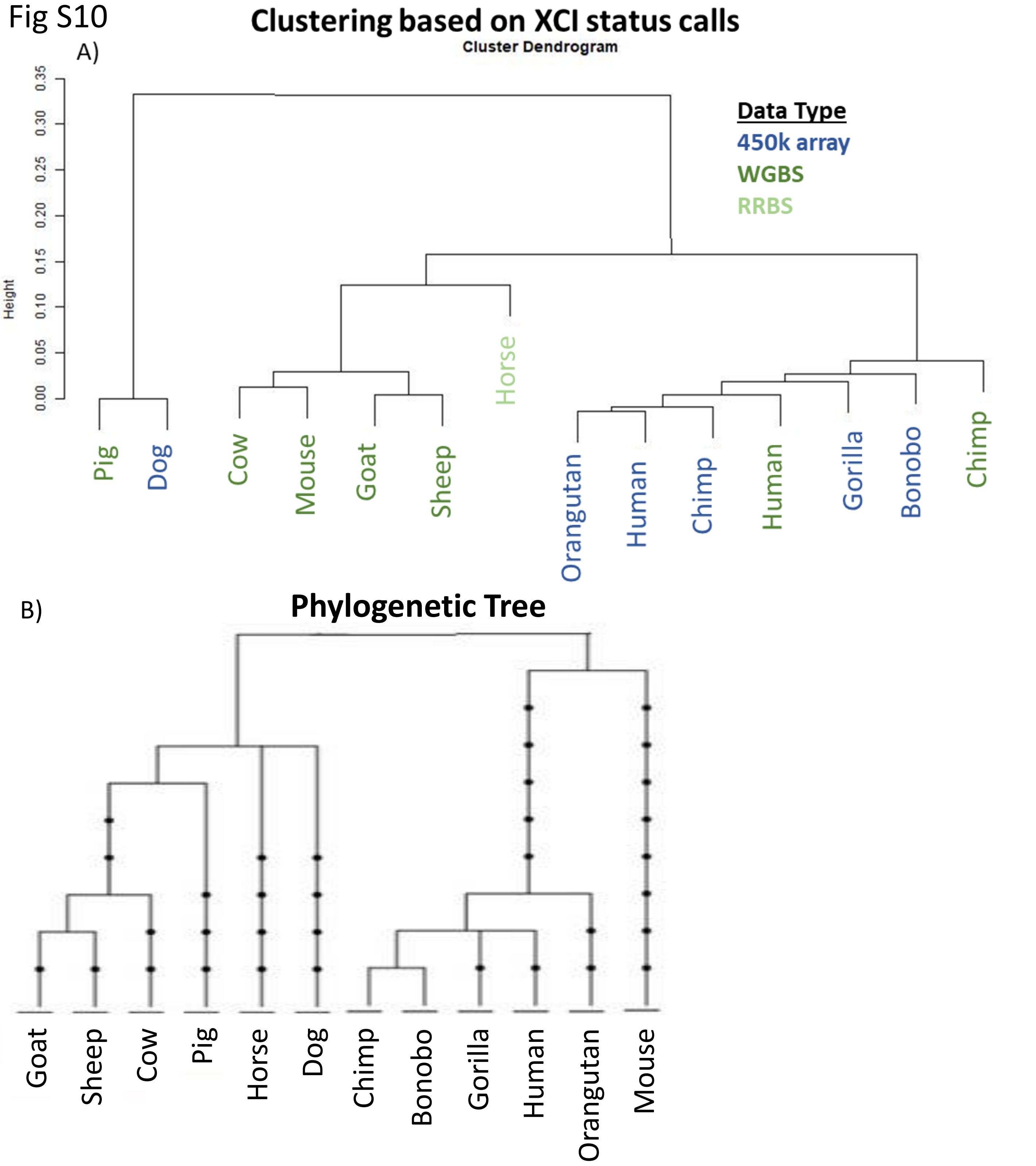


**Figure S10**: **Clustering of species by XCI status calls.** Species were clustered by their XCI status calls (A) and compared to a phylogenetic tree showing their evolutionary relations (B). For the clustering, species names are colored by the type of data used to generate the XCI status calls.
